# LNP Delivery of Prime Editors Enables Durable Correction of PAH R408W in Phenylketonuria Models

**DOI:** 10.64898/2026.10.06.753751

**Authors:** Stephanie Yazinski, Susanna Canali, Yang Xu, Xuling Zhu, James Nelson, Wenguo Gao, Zharko Daniloski, Wendy Leung Owen, Felicity Winkler, Andrea Frassetto, Anna Simon, Jon Penterman, Sean Maguire, Danielle Fuchs, James Rybarski, Tim Towner, Viren Bhatia, Karthik Ramachandran, Karl Anderson, Saakshi Shamanth Donthi, Yogeshwar Kelkar, EunMi Juliana Jung, Xinhua Yan, Lindsey Stretz, Madalee Wulf, Maddie Severson, Boris Reznik, Shannon O’Hayre, Eric Huang, Anthony Forget, Kristine McKinney, Patrick Finn

## Abstract

Phenylketonuria (PKU) is a monogenic metabolic disorder caused by mutations in the phenylalanine hydroxylase (PAH) gene, the most common of which is c.1222C>T (p.R408W). Current treatment options, including dietary restriction and enzyme substitution, can be effective but are often challenging for patients to maintain and do not address the underlying genetic defect. Gene editing represents a compelling one-time treatment approach for PKU. Herein, we demonstrate mRNA:LNP-mediated delivery of prime editors that enable therapeutically-relevant levels of correction of the R408W mutation in both in vitro and in vivo model systems. Further, we found that co-delivery of the DNA repair modulator RAD18 and engineering of the prime editor protein, enabled 2-fold and 3-fold increases in prime editing relative to PE3 respectively, resulting in ∼50% correction efficiency in a humanized PKU mouse model. Collectively, this work identifies novel potential approaches to enhancing prime editing and establishes LNP-delivered prime editing as a mutation-specific, non-viral therapeutic strategy for PKU, as well as providing a framework for extending prime editing to other inherited liver diseases.

## Introduction

Phenylketonuria (PKU) is a rare autosomal recessive metabolic disorder caused by pathogenic mutations in the enzyme phenylalanine hydroxylase (PAH) [1]. PAH is expressed in hepatocytes, where it catalyzes the conversion of phenylalanine to tyrosine. In the absence of functional PAH, phenylalanine accumulates to toxic levels, leading to neurocognitive impairment, developmental delay, and, if left untreated, cumulative brain damage [2]. The most common variant in individuals of European descent is the mutation of Arg408 to tryptophan (p.R408W; c.1222C>T), which significantly reduces PAH stability and activity [3].

Standard management of PKU relies on strict and lifelong dietary restriction of phenylalanine, often combined with synthetic amino acid supplements. While effective in controlling phenylalanine levels, this approach is burdensome and difficult to maintain over time. Two pharmacologic agents, the BH4 cofactor sapropterin dihydrochloride and pegvaliase, an enzyme substitution therapy, are available but have variable efficacy and require ongoing administration [4,5]. These limitations highlight the need for a durable therapy that restores endogenous PAH function.

Genome editing technologies offer a compelling path toward curative treatment of PKU by permanent genomic correction of the underlying genetic defect [6]. Correction of the R408W variant alone would enable treatment of 32.9% of PKU patients in the United States [7]. Multiple potential gene editing approaches could enable treatment of this variant. Insertion of a healthy PAH gene at a safe harbor locus could be enabled either via NHEJ-mediated insertion following a nuclease-generated double-strand break (DSB) or via recombination at a prime editor-installed landing pad, as previously demonstrated [8]. However, DSBs could result in unintended genomic rearrangements, leading to potential genotoxicity. Additionally, the donor gene would likely need to be delivered via a viral vector enabling nuclear import of DNA hepatocytes, whereas other editing systems could be delivered via lipid nanoparticle (LNP) alone. Alternatively, in vivo precision editing could enable correction of the R408W mutation and restore expression of healthy PAH protein. Unlike traditional CRISPR-Cas9 nuclease approaches that rely on double-strand breaks and imprecise repair to generate genomic changes, base editing and prime editing can enable more precise installation of targeted nucleotide changes without requiring donor templates or inducing double-strand DNA breaks [9]. The feasibility of such an approach for the treatment of liver-based rare disease was demonstrated in a landmark 2025 case report, where base editing was successfully used to correct a pathogenic variant in a pediatric patient with a severe urea-cycle disorder [10]. The R408W mutation has been shown to be targetable by both base and prime editors [11,12]. However, while adenine base editors can revert the c.1222C>T mutation, the presence of additional T:A base pairs in the editing window could result in potentially deleterious coding or noncoding variations. Accordingly, we identified prime editing, which can make precise changes to the genome without appreciable bystander editing, as being particularly well-suited for correction of the PAH c.1222C>T variant. Recent advances in LNP delivery have enabled efficient in vivo delivery of RNA-encoded prime editing components, particularly to hepatocytes, which are the principal site of PAH expression [13,14]. These components generally consist of a fusion protein comprising a Cas9 nickase and a reverse transcriptase, as well as a prime editing guide RNA (pegRNA) with an optional nicking guide RNA (ngRNA) [9]. In this study, we developed and tested a prime editing strategy to directly correct the c.1222C>T (p.R408W) mutation in *PAH*, the most common pathogenic variant in European individuals with PKU. As a proof of concept, we employed a pegRNA and ngRNA previously validated by Brooks et al. [12], who demonstrated successful in vivo correction of the R408W variant using an adeno-associated virus (AAV)-delivered prime editor system. Building on this foundation, we formulated mRNA encoding PE2 protein [9] together with the validated pegRNA and ngRNA into LNPs optimized to improve encapsulation and intracellular activity. Using this system, we demonstrated efficient genomic correction of the R408W variant in both Hep3B cells engineered to harbor the mutation and a newly-generated humanized mouse model. Importantly, the level of editing achieved in vivo in our humanized mouse model of R408W-associated PKU was sufficient to restore hepatic PAH expression and reduce blood phenylalanine concentrations below clinically relevant thresholds, constituting a therapeutically meaningful correction. These results establish a non-viral, mutation-specific platform for treating PKU with prime editors.

As well as using previously established prime editing components, we also explored several approaches to enhance correction of the R408W mutation in model systems. Manipulation of DNA repair has been shown to be a powerful lever to improve prime editing efficiency. In addition to the use of ngRNAs [9], inhibition of mismatch repair via installation of synonymous mutations at the target site or supplementation with dominant negative MLH1 protein has been shown to increase the efficiency of point mutation installation via prime editors [15]. Upon screening different DNA repair modulators, we found that addition of mRNA encoding RAD18 - an E3 ubiquitin ligase known for its roles in post-replicative repair, translesion synthesis, and homologous recombination [16, 17, 18, 19, 20] resulted in a two-fold increase in prime editing efficiency in both in vitro and in vivo model systems. Separately, we rationally engineered the prime editor protein to include a D269A point mutation within the Cas9 nickase, previously shown to increase the rate of R-loop formation in the context of the SpRY nuclease [21]. Incorporation of this mutation into a prime editor bearing the previously identified L435K and V223M retroviral point mutations [12] resulted in a 1.5-fold improvement in prime editing efficiency relative to the dual mutant and a 3-fold improvement relative to PE2 in our mouse model.

## Results

### Correction of the PKU-causative R408W mutation in an in vitro model

Evaluation and optimization of gene editing systems in vitro is best done using a relevant cell line model bearing the mutation of interest. To establish such a model for evaluating prime editing of the *PAH* c.1222C>T (p.R408W) mutation, we generated a Hep3B cell line homozygous for the R408W variant via CRISPR-mediated knock-in using a single-stranded DNA oligo donor (Hep3B-R408W). Mutation installation was confirmed by Sanger sequencing of the resulting cells **(Figure 1A).** Consistent with this mutation, we observed a 10-fold reduction in PAH protein in cells bearing this mutation as compared to wild-type Hep3b cells (**Figure 1C**).

**Figure 1:**
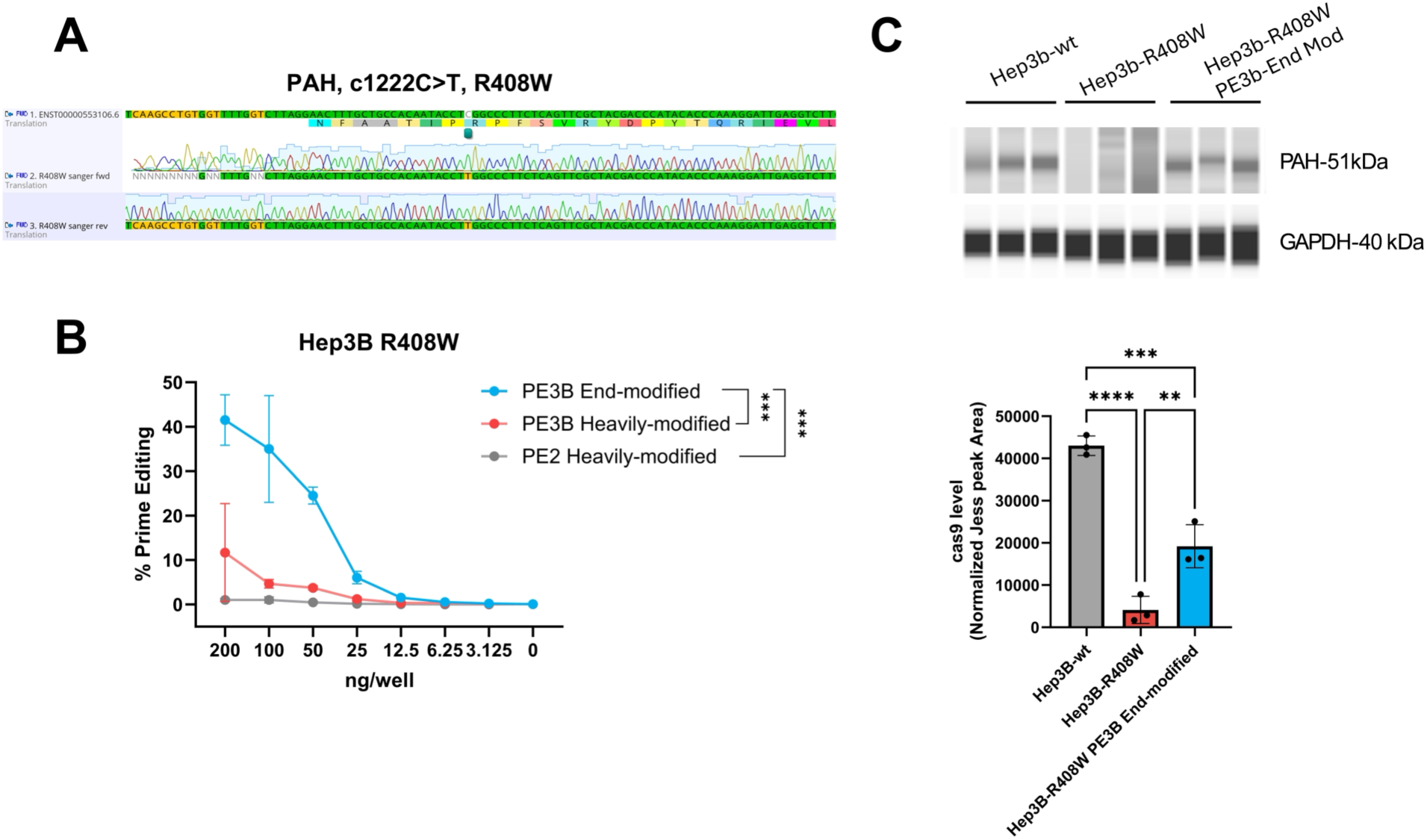
In vitro correction of the PAH R408W mutation in engineered Hep3b cells. (A) Sanger sequencing traces of the PAH locus in engineered Hep3B cells indicating successful knock-in of the c.1222C>T R408W variant. (B) Prime editing correction of the R408W allele in Hep3B-R408W cells using either end-modified (blue circles) or heavily modified guides (red circles), or heavily modified pegRNA without ngRNA (gray circles). Data represent mean ± SD from biological replicates. (C) (top) PAH protein levels as determined by Jess analysis of PAH (51 kDa) and loading control GAPDH (40 kDa) in wild-type Hep3B, Hep3B-R408W, and prime-edited Hep3B-R408W. Densitometric quantification (bottom) indicates a significant decrease in PAH expression in R408W mutant cells and partial restoration after editing. Each dot represents an individual replicate. Statistical significance was assessed by one-way ANOVA followed by Tukey’s multiple comparisons test. Significance levels: p < 0.01 (**), p < 0.001 (***), p < 0.0001 (****).

Previous efforts optimized a pegRNA and PE3b-ngRNA capable of in vitro correction of the R408W mutation in Huh7 cells [12]. This pegRNA templates the R408W correction as well as two additional synonymous mutations (c.1221T>A and c.1224G>T) designed to increase editing efficiency [12], while the ngRNA biases DNA repair in favor of the prime-edited strand. To our knowledge, however, these guides had not been tested in Hep3B cells. To evaluate the ability of these components to affect correction in our model cells, we transfected Hep3B-R408W cells with mRNA encoding the PE2 protein and synthetic pegRNA and ngRNA, delivered via lipoplex. Enhancing guide stability via incorporation of chemical modifications is generally necessary for optimal gene editing efficiency in vivo. We therefore tested versions of these guides with two different scaffold modification patterns: end-modified pegRNA and nicking guides that bear 2’-O-methyl ribose modifications and phosphorothioate linkages at their 5′ and 3′ termini, and heavily-modified guides that contain additional 2’-O-methyl ribose modifications at 40 positions within the tetraloop, direct repeat, and hairpins 1 and 2 [22] **[Table I]**.

As shown in **Figure 1B**, prime editing using the end-modified guides led to robust, dose-dependent correction of the R408W mutation, reaching over 40% editing at the highest dose. Editing in the absence of the ngRNA was negligible. Importantly, we observed minimal bystander edits, scaffold incorporation, or indels as the result of prime editing (<1%; **Supplementary Figures 1A-B).** Editing also led to a partial restoration of wild-type PAH protein and mRNA levels, as measured by Jess capillary Western blot **(Figure 1C).** These protein-level results are consistent with previous observations that the R408W variant reduces PAH stability [23].

In contrast, use of heavily-modified guides yielded substantially reduced editing relative to end-modified guides (∼10%), again with minimal bystander edits, scaffold incorporation, or indels (<1%; **Supplementary Figures 1A-B).** A similar reduction in editing efficiency was observed for a second set of pegRNAs targeting an unrelated site in mouse primary hepatocytes **(Supplementary Figure 1C)**.

Finally, we next assessed the in vitro half-life of the prime editor protein in these cells to better understand the prime editing system (**Supplementary Figure 1D**). The kinetic profile, as measured by Jess using an antibody recognizing the Cas9 portion of the editor, indicates that prime editor protein reaches maximal expression levels around 6-8 hours post-treatment, is sustained through hours 8-16, and gradually declines thereafter. Together, these results demonstrate that the engineered Hep3B-R408W cell line, paired with published pegRNA designs and optimized scaffold chemistries, provides a useful platform for evaluating prime editing efficiency, precision, and functional outcomes in the context of the R408W PKU mutation. We expanded this approach by generating other cell lines each bearing different pathogenic, PKU-causative mutations (R261Q (c.782G>A) and R243Q (c.728G>A)), as well as additional mutations that result in other monogenic liver diseases (*CBS* I278T (c.833T>C) and *G6PC* Q347* (c.1039C>T)). In each case, we designed and tested pegRNAs and ngRNAs to correct these mutations, with prime editing efficiencies ranging from 5 to 30%, depending on the target mutation (**Supplementary Figure 2-3**).

### Chemical modification of the scaffold of the R408W-targeted pegRNA reduces prime editing efficiency

We hypothesized that the incorporation of chemical modifications in the pegRNA scaffold could reduce prime editing efficiency by inhibiting binding of the prime editor protein to the pegRNA or binding of the prime editor RNP to the genome. To investigate these possibilities, we adapted an in vitro FRET-based assay to measure R-loop formation [24]. In this assay, double-stranded DNA corresponding to the R408W target site is modified with a Cy3 fluorophore on the non-target strand two nucleotides upstream of the cleavage site and Cy5 fluorophore on the target strand 6 nt downstream of the cleavage site. R-loop formation increases the distance between the two fluorophores, decreasing the observed FRET signal. We validated this approach by testing a version of the guide RNA bearing a mutated scaffold that inhibits DNA cleavage, and observed a meaningful reduction in R-loop formation (**Supplementary Figure 6**).

We measured the decrease in FRET over time after adding catalytically-inactive Cas9 protein to a solution containing guide RNA + target DNA. We tested end-modified and heavily modified sgRNAs without a prime editing extension to prevent potential confounding effects due to spacer: PBS hybridization. Upon doing so, we observed near identical rates of R-loop formation with both guides, suggesting that, in the absence of the pegRNA extension, the heavily modified guide did not reduce the rate of prime editor binding or R-loop formation. Similarly, the heavily modified and end-modified sgRNAs showed near identical cleavage kinetics of target DNA in a cleavage assay using catalytically-active Cas9 protein, suggesting that the discrepancy in activity in cells did not arise from differences readily detected by these biochemical assays.

Next, we sought to determine the regions of the pegRNA scaffold where chemical modification resulted in reduced prime editing efficiency. To do so, we tested pegRNAs bearing different combinations of 2′OMe modifications within the direct repeat, tetraloop, and hairpin 1 and 2 regions of the guide scaffold. In PHH cells, retention of the 2′-OMe modification in hairpin 2 and the tetraloop was sufficient to reduce activity, whereas modification confined to the direct repeat region was well-tolerated (**Supplementary Figure 4**). However, since guide stability requirements are significantly more stringent in vivo than in vitro, we suspected only modifying the direct repeat would not significantly improve prime editing relative to end-modified guides. Accordingly, for subsequent in vivo experiments we decided to test both end- and heavily-modified pegRNAs.

In addition to testing modified the above scaffold modification patterns and sequences, we tested several additional variations of the PAH R408W pegRNA (**Supplementary Text**). While these studies are limited, they suggest that chemical modification of the PBS and genome-matched scaffold sequences may improve pegRNA performance. However, we speculate that the ability to tolerate chemical modification of the pegRNA scaffold is likely to be pegRNA sequence and genomic site dependent.

### In vivo correction of *PAH R408W* using LNP-delivered Prime Editing in a published PKU mouse model

Most mouse models of PKU, such as ENU2, do not contain the humanized *PAH* gene or mutations relevant to human disease. Therefore, to evaluate the therapeutic potential of our prime editing strategy in vivo, we first utilized a previously characterized *Pah^R408W^* knock-in mouse model of phenylketonuria (PKU), generated at the University of Pennsylvania (*Pah*^R408W^-UP) [12]. This model accurately recapitulates key biochemical and molecular hallmarks of human disease [12]. As shown in **Supplementary Figure 7A**, wild-type mice of both sexes exhibit low circulating phenylalanine (Phe) concentrations, while animals that are heterozygous for the R408W mutation display a moderate elevation in Phe levels. In contrast, homozygous *Pah*^R408W^-UP and ENU2 knock-in mice lines demonstrate markedly elevated Phe levels, with ENU2-Hom animals exceeding 3000 µM, consistent with a severe PKU phenotype.

Liver PAH protein expression, quantified by mass spectrometry, mirrored these biochemical findings (**Supplementary Figure 7B)**: wild-type and heterozygous animals showed robust hepatic PAH expression, whereas levels were minimal to undetectable in homozygous *Pah^R408W^*-UP mice. These data confirm the expected genotype-phenotype relationship in both mouse models and establish a validated platform for in vivo testing of prime editing-mediated phenotypic correction.

After validating this disease model, we set out to determine whether LNP-mediated delivery of prime editor mRNA and pegRNA could achieve functional correction at the *PAH p.R408W* site. In our initial experiment, we formulated lipid nanoparticles containing i) mRNA encoding PE2 protein, ii) heavily modified ngRNA, and iii) either end-modified or heavily modified pegRNA **(Table I),** delivering them to mice that were homozygous or heterozygous for the p.R408W mutation. Although we had observed a decrease in prime editing efficiency in vitro with scaffold-modified pegRNAs, we speculated that their improved stability might yield better prime editing in vivo. Mice received a single intravenous dose of 3 mg/kg LNP-formulated with mRNA and guide RNAs, at a ratio of 4:3:1 mRNA: pegRNA:ngRNA. To confirm our ability to edit this site with LNP-delivered gene editing components, we also formulated LNPs bearing mRNA encoding wild-type Cas9 and chemically-modified sgRNA targeting the p.R408W locus. Liver tissue was harvested one-week post-injection for analysis of editing efficiency and phenylalanine levels. We observed significant DSB formation in mice treated with wild-type Cas9 targeted to *PAH*, confirming the ability to edit at this locus in the mouse model. Targeted NGS confirmed editing at the *PAH* locus in mice treated with prime editors, with correction rates up to 10% observed in whole liver **(Figure 2A).** This molecular correction translated into a substantial and sustained reduction in plasma phenylalanine concentrations **(Figure 2B).** PAH protein expression was restored in edited livers, as assessed by immunoblot and immunohistochemistry **(Figures 2C).** Prime editing at an unrelated locus (Dnmt1) had no effect on Phe or PAH levels. Interestingly, we found that use of either end- or heavily-modified pegRNA resulted in similar levels of prime editing efficiency in vivo, suggesting that the likely enhanced stability of the heavily-modified pegRNA can compensate for any reduction in enzymatic activity. Importantly, we observed negligible indels or scaffold incorporation at the target site following prime editing **(Supplementary Figure 8).** These results demonstrate that LNP-delivered prime editing can efficiently correct the R408W mutation in vivo and restore metabolic function in a published humanized PKU model.

**Figure 2:**
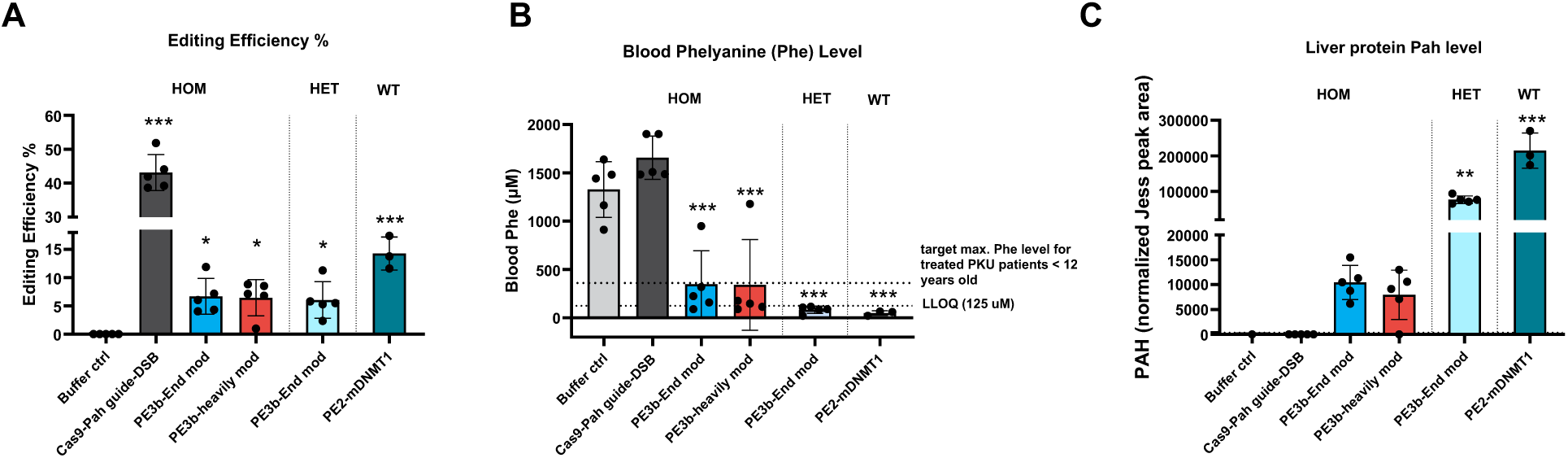
Assessment of editing efficiency, blood phenylalanine levels, and hepatic PAH protein restoration following prime editing treatment in PKU mice. Homozygous (HOM), heterozygous (HET), and wild-type (WT) mice from the University of Pennsylvania were treated with 3 mg/kg of LNPs containing prime editor components. Mice were treated with mRNA encoding PE2 and pegRNA and ngRNA targeting the PAH locus. Additional, control groups were treated with either i) buffer alone, ii) mRNA encoding wtCas9 and sgRNA targeting the PAH locus or iii) mRNA encoding PE2 and pegRNA targeting the DNMT1 locus. (A) Editing efficiency at the PAH locus in liver tissue, as measured by targeted next-generation sequencing one-week post-treatment. (B) Blood phenylalanine (Phe) levels quantified by LC-MS/MS one-week post-treatment. Dashed lines indicate the target maximum Phe concentration (360 µM) for treated PKU patients <12 years old [25] and the lower limit of quantification (LLOQ, 125 µM). (C) PAH protein levels in liver lysates measured by capillary-based immunodetection (Jess, ProteinSimple), reported as normalized peak area. Data are presented as mean ± SD. Each point represents an individual animal. Statistical significance was assessed by one-way ANOVA followed by Dunnett’s multiple comparisons test versus the buffer-treated control group. Significance levels: p < 0.05 (*), p < 0.01 (**), p < 0.001 (***).

### Generation and characterization of a humanized mouse model carrying the PAH R408W Mutation

To facilitate further in vivo evaluation of prime editing, we generated our own humanized knock-in mouse model carrying the *PAH* R408W mutation in collaboration with Taconic (*Pah^R408W^*). This model was developed using CRISPR/Cas9 technology to insert the human PAH R408W mutation into the C57BL/6NTac mouse background. Similar to the University of Pennsylvania mice, this strategy enabled partial humanization of the *Pah* gene through CRISPR/Cas9-mediated gene editing. Specifically, a region spanning 50 bp upstream to 50 bp downstream of the R408 codon in exon 12 was replaced with the corresponding human genomic sequence. The Cas9 protein, guide RNAs (gRNAs), and donor oligonucleotides were microinjected into C57BL/6NTac zygotes to achieve the targeted integration **[Supplementary Figure 9**]. As a control for this disease animal model, we employed the same strategy to generate mice with the same humanized PAH gene region but without the R408W point mutation (*Pah^R408^*). These animals are expected to express a chimeric protein without exhibiting a disease phenotype, maintaining normal Phe levels. This approach allows us to assess gene expression in the absence of disease mutation.

To validate the phenotype of our PKU mouse model, we first measured blood Phe levels and PAH protein expression. As expected, female homozygous (Hom) *Pah^R408W^* mice showed elevated Phe levels, comparable to those seen in ENU2 mice, confirming the disease phenotype **(Figure 3A).** These animals also displayed reduced PAH protein levels, consistent with a loss-of-function allele **(Figure 3B)**, effectively recapitulating the human disease phenotype.

**Figure 3:**
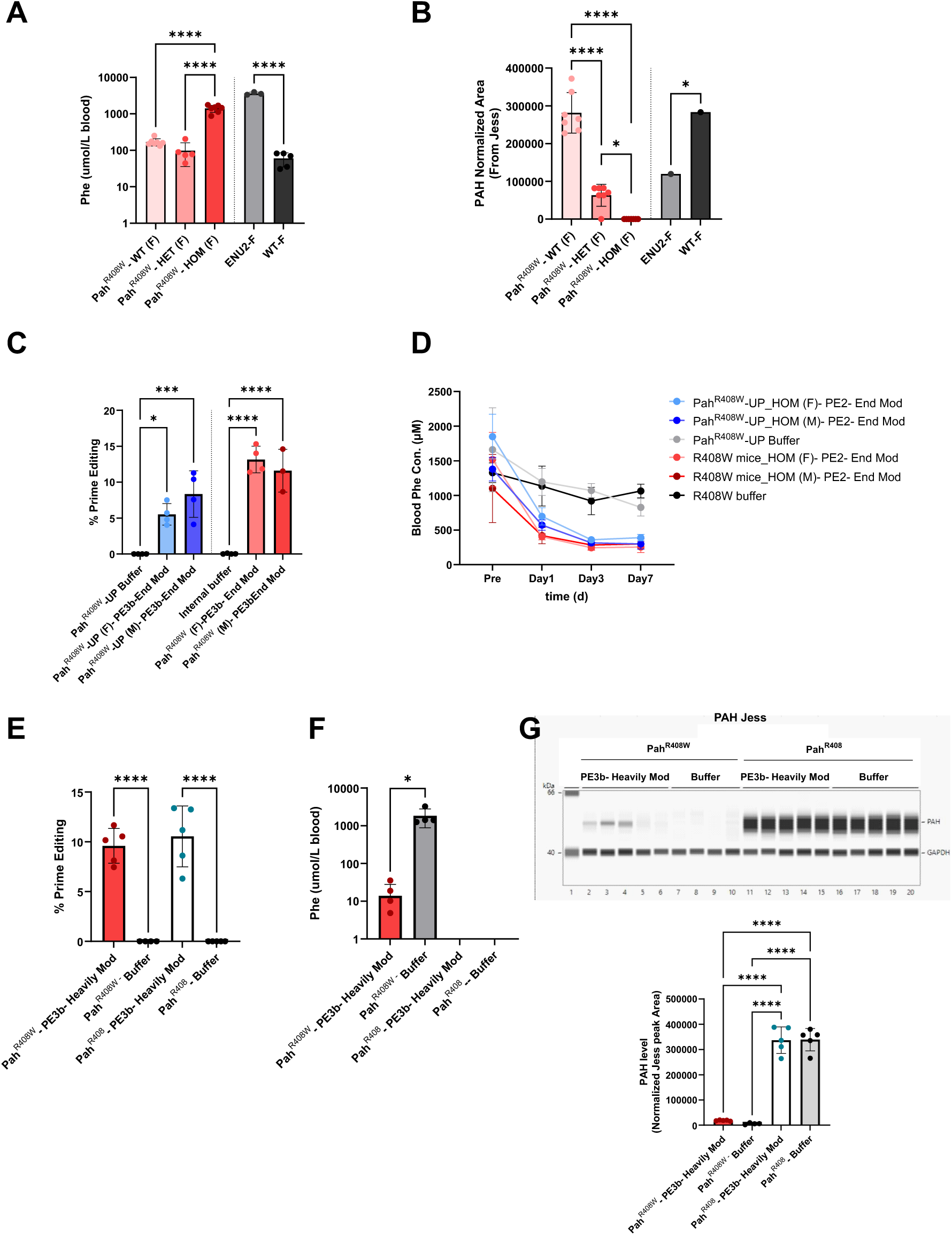
Editing comparison between the published in vivo animal model Pah^R408W^-UP and our internal transgenic knock-in Pah^R408W^. (A-B) Baseline phenylalanine (Phe) levels and hepatic PAH protein expression in male and female Pah^R408W^ mice compared to heterozygous, ENU2, and wild-type controls. (A) Blood Phe concentration was measured by LC-MS/MS and reported as µmol/L. (B) Liver PAH protein levels were quantified by Jess and reported as normalized peak area. (C-D) Correction of the Pah R408W mutation by LNP-delivered PE2 using end modified pegRNA in Pah^R408W^ and PahR408W-UP mice. (C) Prime editing efficiency in liver was assessed by targeted next generation sequencing seven days post-treatment. (D) Blood Phe concentrations were measured by LC-MS/MS at baseline and one, three, and seven days post-treatments. All compared samples were collected, stored, and measured together. (E-F) Additional in vivo assessment of prime editing and Phe levels following prime editing with heavily modified pegRNA, measured identically to as described in (C-D) above. (G) PAH protein levels in liver lysates were analyzed by Jess. (top) Representative image showing PAH and GAPDH detection. (bottom) Quantification of PAH expression reported as normalized peak area. Data are presented as mean ± SD. Each point represents an individual animal. Statistical significance was assessed by one-way ANOVA followed by Dunnett’s multiple comparisons test versus the buffer-treated control group. Significance levels: p < 0.05 (*), p < 0.01 (**), p < 0.001 (***), p < 0.0001 (****).

After confirming the expected biochemical phenotype, we compared prime editing efficiency in our internal R408W (*Pah^R408W^*) mouse model with the UPenn model previously described (*Pah^R408W^*-UP) **(Supplementary Figure 7).** Using the previously published pegRNA and nicking sgRNA [12], we achieved comparable editing levels, with up to 15% prime editing observed in our internal model without significant indel formation or scaffold incorporation (<0.2%) (**Figure 3C** and **Supplementary Figure 10**).Consistent with those data, a single administration of prime editor mRNA and guide RNAs resulted in a rapid reduction of Phe levels, reaching a near-therapeutic range by Day 1, with the effect sustained through Day 7 post-injection (**Figure 3D**).

To further characterize the internal model, we compared editing outcomes in R408W mutant mice and control mice that carry only the humanized PAH gene without the mutation. We speculated that mice without the mutation might exhibit lower prime editing efficiency relative to mice with the mutation, since the resulting prime edit has fewer changes relative to the target sequence and thus might be more likely to be repaired by mismatch repair [15]. However, both genotypes showed similar levels of prime editing (**Figure 3E).** Consistent with previous data, elevated baseline Phe levels were observed only in the R408W, which significantly decreased following treatment **(Figure 3F).** Finally, one week following treatment, we observed a small increase in PAH expression in edited *Pah^R408W^*mice, relative to unedited mice (**Figure 3G**).

### Prime editing enables durable expression of PAH and reduction of Phe levels

A significant advantage of gene editing is that a single treatment should enable a durable change in phenotype. To confirm that correction of the p.R408W mutation resulted in durable correction of the PKU phenotype in mice, we designed an in vivo study to monitor the levels of PAH protein and Phe over time and compared it to mRNA-based PAH replacement therapy. *Pah^R408W^* mice were treated either with a single dose of LNP-prime editor or of an mRNA encoding for wt PAH protein (mRNA-PKU), and Phe was measured at 6 hours, 24 hours, or seven days post dose. Importantly, we observed similar levels of prime editing in mice dosed with 3 mg/kg LNP seven days following treatment compared to our previous experiment (**Figure 4A**). Additionally, we observed a sustained reduction in blood Phe beginning at 24 hours post-injection in prime edited mice, which was maintained through Day 7. In contrast, animals treated with mRNA-PKU showed an earlier but transient decrease in Phe levels detectable at 6 hours post-injection, with a rebound to baseline by Day 7 (**Figure 4B**). Buffer-treated *Pah^R408W^*mice and wild-type controls remained unchanged. These findings confirm that prime editing induces a more durable biochemical correction than transient mRNA-based enzyme replacement in this model.

**Figure 4:**
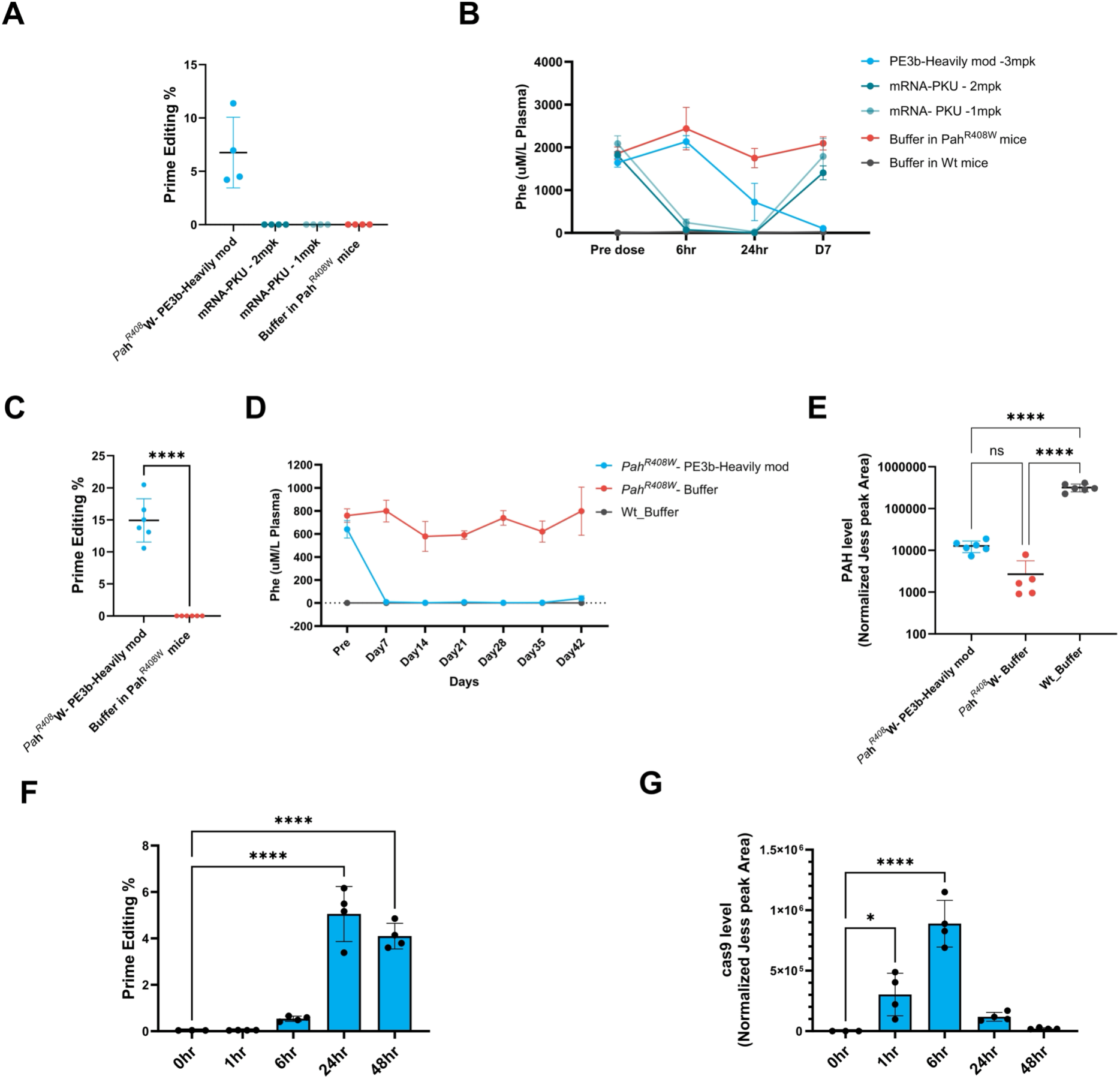
Comparison of phenylalanine correction kinetics and molecular dynamics following prime editing or mRNA-based PAH replacement in PKU mice. (A) Prime editing efficiency in Pah^R408W^ mice treated with PE3b-Heavily mod (3 mg/kg) compared to animals treated with mRNA encoding PAH (1 or 2 mg/kg), and respective buffer controls. Editing was quantified in liver tissue at day 7 post-injection by targeted amplicon sequencing. (B) Plasma phenylalanine (Phe) concentrations were measured at baseline (Pre dose) and at 6 hours, 24 hours, and 7 days post-treatment in Pah^R408W^ homozygous mice. Animals were treated with either LNP-formulated PE3b-Heavily mod prime editor (3 mg/kg) or mRNA encoding PAH at 1 mg/kg or 2 mg/kg. Buffer-treated Pah ^R408W^ mice and wild-type mice served as controls. (C) Prime editing efficiency in the PahR408W mice treated with PE3b-Heavily mod (3 mg/kg) and respective buffer control for the durability study. Editing was quantified in liver tissue at day 42 post-injection by targeted amplicon sequencing (D) Longitudinal monitoring of plasma Phe levels in Pah^R408W^ mice treated with PE3b-Heavily mod (3 mg/kg) or buffer control, compared to wild-type mice, up to 42 days post-treatment. (E) Hepatic PAH protein levels measured at the take down at day 42 post-treatment by capillary electrophoresis (Jess) in Pah^R408W^ mice treated with PE3b-Heavily mod or buffer, and in buffer-treated wild-type mice. (F) Editing kinetics were evaluated in Pah ^R408W^ mice treated with LNP-formulated PE3b-Heavily mod by collecting liver tissue at 1-, 6-, 24-, and 48-hours post-injection. Prime editing efficiency was quantified by targeted amplicon sequencing and reported as percent intended edit. (G) PE2 protein expression in the liver was assessed by capillary electrophoresis (Jess, ProteinSimple) at the same time points. Data are presented as mean ± SD. Each point represents an individual animal. Statistical significance was assessed by one-way ANOVA followed by Tukey’ s multiple comparison test (E) or Dunnett’s multiple comparisons test versus the control group (F, G). Significance levels: p < 0.05 (*), p < 0.0001 (****), ns = not significant.

To assess long-term durability, we followed a separate cohort of *Pah^R408W^* mice out to 42 days post-treatment **(Figure 4C-D).** Again, *Pah^R408W^* mice were dosed once either with LNP-prime editor or buffer as a control. Prime editing reached 20% with a mean of 15% at the 42 day end point. Plasma Phe levels remained stably reduced in edited animals, whereas buffer-treated *Pah^R408W^* mice maintained elevated levels throughout the study. Wild-type animals remained unaffected. At Day 42, liver tissue was collected to quantify PAH protein, where a visible but statistically insignificant increase in hepatic PAH levels in edited was observed **(Figure 4E).**

Finally, to investigate the molecular kinetics of editing, we conducted a separate experiment where we monitored editing and protein levels for two days following dosing **(Figures 4F-G).** Liver samples were collected at 1, 6, 24, and 48 hours post-injection of LNPs containing mRNA encoding PE2, heavily-modified pegRNA, and ngRNA. Targeted sequencing revealed that editing was first detectable at 6 hours, peaked at 24 hours, and remained high through 48 hours **(Figure 4F).** In parallel, PE2 protein levels peaked at 6 hours and declined sharply by 24-48 hours **(Figure 4G**), consistent with rapid translation and clearance of the LNP-delivered mRNA payload.

Together, these results demonstrate that a single administration of LNP-formulated prime editing reagents induces durable gene correction and biochemical rescue in a mouse model of PKU.

### Enhancement of prime editing correction through manipulation of DNA repair RAD18 improves Prime editing

Most gene editing modalities take advantage of one or more strategies to bias cellular DNA repair machinery in favor of incorporating the desired edit. Use of a secondary, nicking guide or incorporation of silent, non-coding edits has been shown to improve prime editing efficiency via evasion of mismatch repair pathways, as has simultaneous delivery of dominant-negative MLH1 protein [15]. However, since different cell types express different DNA repair pathways, this approach is inherently cell-type dependent.

We sought to identify DNA repair factors that could enhance prime editing efficiency or specificity in primary human hepatocytes. To do so, we pursued several experimental paths in parallel. First, we screened a library of known small molecule DNA repair agonists and antagonists to identify compounds which affect prime editing efficiency. Second, we rationally identified a panel of possible DNA repair proteins that if overexpressed could manipulate prime editing outcomes and transfected mRNA encoding them into PHH cells. Third, we performed a lentiviral-based CRISPRi knockout screen in PHH, monitoring prime editing outcomes at a lentivirally-integrated target site.

The mechanism of prime editing involves the creation of nicks, 3’ and 5’ flap structures, and homologous recombination-like intermediates. Accordingly, we hypothesized that RAD18, an E3 ubiquitin ligase known for its roles in post-replicative repair, translesion synthesis, and homologous recombination, [17, 18, 19, 20] might enhance prime editing. We compared the effect of co-expression of either RAD18 or GFP with PE2 across a variety of effector to editor ratios on the efficiency of prime editing at the p.R408W locus. As shown in **Figure 5**, co-expression of RAD18 resulted in significant increases in prime editing efficiency across nearly all ratios of PE2 to RAD18, compared to GFP. This increase was observed across both primary human hepatocytes (PHH) **(Figure 5A** and **5B)** and primary mouse hepatocytes (PMH) derived from *Pah^R408W^* mice **(Figure 5C** and **5D****) and** was consistent across two stoichiometric ratios between the prime editor and the guides (4:3:1 and 8:4:1 PE2, to pegRNA to nicking gRNA, respectively). In PHH, RAD18 co-delivery increased editing from ∼10% to over 20% at optimal dosing. In primary mouse hepatocytes from the *Pah^R408W^* mouse model, RAD18 raised editing from ∼8% to >15%, and in some cases, as high as 30-35%. The effect was dose-dependent, with peak enhancement observed between 100-150 ng total RNA input. Importantly, this increase in prime editing did not lead to any notable increase in indels or scaffold incorporation (**Supplementary Figure 11).** Given its mechanism of action in promoting DNA synthesis at stalled forks and stabilizing editing intermediates, [17, 18, 19, 20, 26] RAD18 may be especially well-suited to support prime editing in non-dividing or post-mitotic cells, such as hepatocytes. However, further research is needed to determine to what extent RAD18 addition can improve prime editing in other cell types and at other genomic loci. Additionally, given RAD18’s role in regulating DNA repair pathways, and reports that RAD18 overexpression can promote unregulated translesion synthesis (TLS), it will be important to assess potential guide-independent genomic alterations or increased mutagenesis [20, 27].

**Figure 5:**
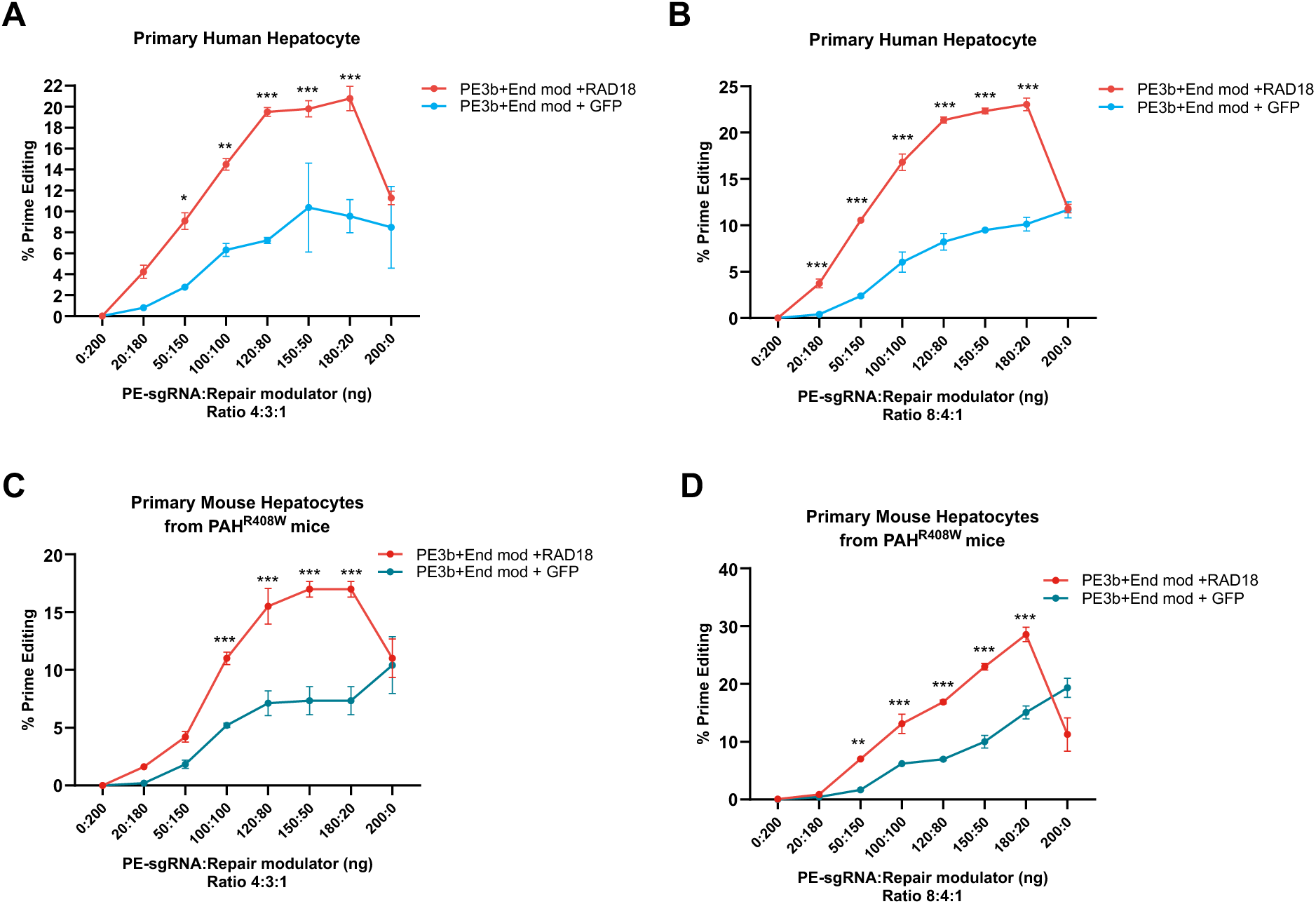
RAD18 enhances prime editing efficiency in PHH and PMH isolated from transgenic PKU^R408W^ mice. (A-B) Prime editing (PE) efficiency at the p.R408W locus in PHH transfected with PE2, pegRNA, ngRNA and either RAD18 or GFP. The total amount of editing reagents was kept constant while the ratio of RAD18 or GFP to prime editor components was varied. Prime editing components were transfected at a ratio of 4:3:1 PE2:pegRNA:ngRNA (A) or 8:4:1 (B). (C-D) Prime editing efficiency in primary mouse hepatocytes derived from Pah^R408W^ mice, as in A and B. Data represent mean ± SD. Statistical significance was assessed by two-way ANOVA, followed by Dunnett’s or Sidak’s multiple comparisons test, as appropriate, comparing each concentration point to the GFP control group. Significance levels: p < 0.05 (*), p < 0.01 (**), p < 0.001 (***).

### Installation of a Cas9 mutation known to speed R-loop resolution increases prime editing

Unlike Cas9 nucleases, the prime editor must maintain DNA binding while coordinating pegRNA pairing and reverse transcription, a structurally complex and potentially unstable intermediate [28]. We speculated that mutations that help to stabilize this intermediate or speed up its formation could improve prime editing efficiency. Further, since extension of the DNA substrate by the RT might result in disruption of the pegRNA DNA duplex and intermediate destabilizations [28], such mutations could also potentially enable the use of longer reverse transcription templates (RTTs), increasing the genomic window available to prime editors. Recent work has shown that removing the physical barrier to R-loop propagation via installation of a D269A mutation in SpRY can greatly increase the rate of R-loop formation at target strand cleavage [21]. We hypothesized that accelerating the formation of the prime editing intermediate and cleavage of the target strand via incorporation of this mutation could improve editing efficiency.

To test this approach, we designed versions of SpCas9-MMLV RT fusion proteins bearing different mutations designed to improve binding to the genomic site and examined their prime editing efficiency. We initially designed constructs that bore either no additional mutations relative to the previously used prime editor system (PE2), a pair of mutations previously reported to improve the solubility and binding of the RT to dNTPs (V223M and L435K, numbering relative to the RT sequence alone, PE2-KM) [29]; or these two mutations and the D269A mutation (PE2-K_M_-D269A) **(Table VI).** We transfected mRNAs that encode these editors, along with the pegRNA and ngRNA used to correct the PAH R408W mutation into PHHs **(Figure 6A).** Consistent with previous findings, [21, 29] we observed a significant increase in editing efficiency with PE2-K_M_ compared to PE2. Interestingly, addition of the D269A mutation to PE2-K_M_ resulted in a significant increase in prime editing efficiency.

**Figure 6:**
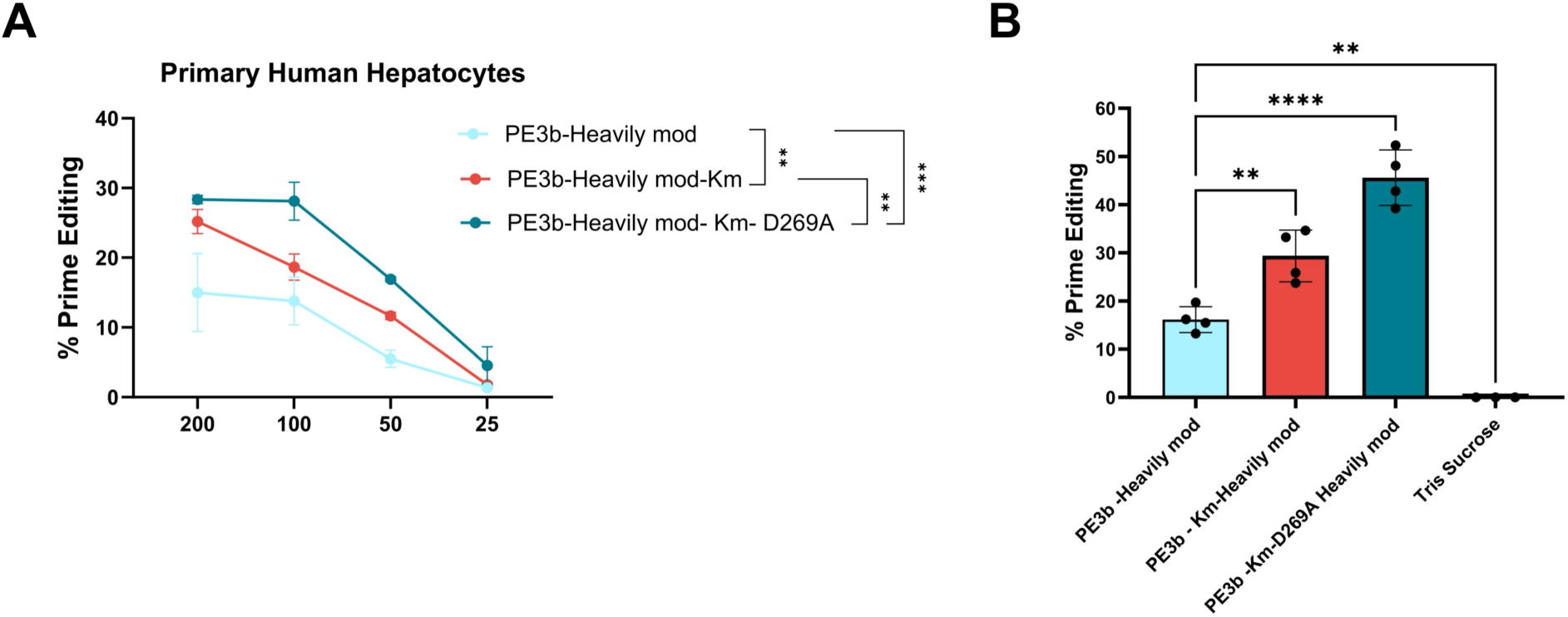
Incorporation of a D269A mutation into the Cas9 nickase enhances prime editing efficiency in human hepatocytes and in vivo. (A) Primary human hepatocytes (PHH) were transfected with mRNA encoding PE2 variants: PE2, PE2-KM (containing RT mutations V223M and L435K), or PE2-KM-D269A. (B) In vivo prime editing in PahR408W mice treated with a single intravenous dose (3 mg/kg) of LNP formulated with the indicated prime editor mRNA and heavily modified modified pegRNA and ngRNA. Mice were sacrificed one-week post-treatment, and editing efficiency in liver tissue was quantified by targeted next-generation sequencing. Data are presented as mean ± SD. Each point represents an individual animal. Statistical significance was assessed by one-way ANOVA followed by Dunnett’s multiple comparisons test versus the PE3b Heavily mod group. p < 0.01 (**), p < 0.001 (***), p < 0.0001 (****)

In vivo evaluation of PE2-K_M_-D269A also showed a clear boost in performance relative to PE2 or PE2-K_M_ **(Figure 6B).** While the baseline PE2 construct achieved only ∼15% prime-edited alleles and PE2-K_M_ yielded about 30%, PE2-K_M_-D269A yielded∼45% prime editing efficiency, a roughly three-fold improvement over the original PE2 and a ∼50 % gain relative to the K_M_ variant alone. No significant increase in indel formation or scaffold incorporation was observed in animals treated with PE2-K_M_ or PE2-K_M_-D269A relative to animals treated with PE2 **(Supplementary Figure 12).** While encouraging, further work is needed to determine whether the D269A prime editor variant results in improved editing in other cell types or at other genomic loci.

## Discussion

This study demonstrates that lipid nanoparticle (LNP)-delivered prime editing can durably and precisely correct the PAH p.R408W mutation in cellular and animal models of phenylketonuria (PKU), offering a promising non-viral, single-dose genetic therapy for this lifelong metabolic disorder.

We engineered a Hep3B cell line to bear the p.R408W mutation and evaluated a previously identified prime editing system [12] for its ability to correct this mutation, achieving up to ∼50% correction in Hep3B cells engineered to bear the p.R408W mutation. These edits were accompanied by partial restoration of PAH protein levels, with minimal undesired on-target effects such as indels or scaffold incorporation, key features for therapeutic safety. While heavily modified guides likely offered improved chemical stability, they showed reduced editing efficiency in vitro, highlighting the importance of balancing chemical protection with functional activity.

To demonstrate the ability of this platform to effect gene correction in vivo, we leveraged two complementary mouse models of PKU: the established *Pah^R408W^*-UP strain from the University of Pennsylvania [12] and a newly developed humanized knock-in model generated via CRISPR/Cas9 (*Pah^R408W^*). Both recapitulate hallmark features of PKU, including elevated phenylalanine (Phe) levels and reduced hepatic PAH expression. Following a single 3 mg/kg intravenous dose of LNP-formulated PE2-encoding mRNA and guides, we observed therapeutic correction in both models: prime editing rates up to 15% in liver, robust reduction of plasma Phe to near-physiological levels, and increased PAH protein expression. These results represent a durable phenotypic rescue of PKU via RNA-based delivery of a prime editor.

Importantly, our study demonstrates that LNP-mediated delivery of the exact same pegRNA and nicking guide sequences used in a prior AAV-based study [12] can achieve therapeutic correction in vivo. While the AAV-delivered system required viral vectors to achieve sufficient editing, our LNP approach matched or exceeded editing performance without viral vectors, offering a more transient, re-dosable, and potentially safer modality. This result addresses a key translational bottleneck previously identified in the field-namely, the challenge of efficiently delivering prime editing components using non-viral delivery systems and establishes that LNPs are a viable and clinically scalable platform for prime editing of hepatic targets.

We compared prime editing with mRNA-based enzyme replacement using synthetic PAH mRNA. While both approaches lowered plasma Phe, the mRNA-based effect was short-lived, peaking at 6 hours and returning to baseline by Day 7. In contrast, LNP-delivered prime editing induced a delayed but durable correction, with effects sustained over one week. Kinetic profiling showed that Cas9 expression peaked at 6 hours and editing efficiency at 24 hours, consistent with rapid translation of the LNP-delivered mRNA and accumulation of genomic edits thereafter. These findings underscore the longitudinal advantage of genome editing over transient gene supplementation, even with a single dose.

We also explored ways to further improve prime editing efficiency. Overexpression of the DNA repair factor RAD18 consistently enhanced editing in both human and mouse primary hepatocytes, increasing editing levels by 2-3-fold. This aligns with RAD18’s known role in post-replicative DNA repair and supports its utility as a co-factor for prime editing, particularly in non-dividing cells like hepatocytes. Similarly, installation of the D269A mutation in the Cas9 domain of the prime editor, previously shown to accelerate R-loop formation in an expanded PAM Cas nuclease [21] resulted in ∼50% higher editing versus the corresponding PE2-K_M_ variant. Together, these findings point to new avenues for improving editing efficiency through engineering of the editor or modulation of cellular repair pathways, especially in clinically relevant tissues. However, given the limited scope of cell types, genomic loci, and installed mutations examined here, further testing is needed to determine how broadly applicable to prime editing these approaches are.

Finally, extensive biochemical and structural work informed our design choices. Our data support that end-modified guides do not impair R-loop formation or Cas9 cleavage, as confirmed by FRET and cleavage assays. Further pegRNA engineering experiments targeting the 3′ extension, scaffold sequence, and spacer length underscore the critical and sometimes underappreciated role of pegRNA structure-function relationships in determining prime editing outcomes. Optimization of these components, including use of genome-matched scaffolds and chemically modified PBS sequences, allowed us to fine-tune activity and minimize scaffold incorporation, a potential safety concern.

This work arrives at a time of rapid progress in therapeutic genome editing, where proof-of-concept studies are rapidly translating into clinical interventions. In a landmark milestone this year, the first in vivo gene editing treatment in a pediatric patient with a rare urea cycle disorder was successfully reported [10]. That achievement highlights the real-world feasibility of using precision gene editing to treat monogenic liver diseases. Our findings build on this momentum, showing that prime editing delivered via LNPs, without the use of viral vectors, can restore gene function and correct disease phenotypes with high precision and therapeutic impact.

In conclusion, this study establishes that LNP-delivered prime editing is a viable therapeutic strategy for PKU, capable of achieving single-dose, durable gene correction and phenotypic rescue in vivo. The integration of repair pathway modulation, editor protein engineering, and pegRNA design optimization offers a blueprint for extending prime editing into other monogenic liver diseases. As LNP and prime editing technologies continue to mature, this platform may pave the way for mutation-specific, non-viral genome editing therapies in clinical settings.

## Material and Method

### mRNA-LNP production and formulation

Each mRNA and guide were coformulated in LNPs containing an ionizable lipid, distearoylphosphatidylcholine, cholesterol, and PEG2k-DMG lipids, and administered intravenously. mRNA constructs were manufactured *in vitro* via T7 RNA polymerase-mediated transcription with complete replacement of uridine by N1-methyl-pseudouridine, while guides were procured from an external CRO. [25].

### Editing quantification

Genomic DNA from hepatocytes or mouse tissue was extracted as described below and the human locus encompassing the sgRNA target site amplified via PCR using primers flanking the target site (AGAGTCAGCATGCCAATATGTGT and CGATGGTAGGGAAAGACAGTCTTCGATTA). The chimeric PAH locus in the engineered mouse-material was amplified using primers CACAGCAAGGAAAAAGCCACTTGGAA and CCCAGCAGGTGTGAGCTACTTACTATTAA. Each primer set had appended Illumina handles on the 5’ end. Resulting amplicons were barcoded, pooled, and sequenced on an Illumina instrument to quantify editing rates. Amplicon sequencing data were quantified using CRISPResso2.0 or an equivalent pipeline.

### In Vitro Transfection of Primary Human Hepatocytes

Primary Human hepatocytes (BioIVT) were seeded at 40,000 cells/well in collagen-coated 96-well plates using proprietary hepatocyte plating medium. After 4 hours, the medium was replaced with hepatocyte maintenance medium supplemented with 5% FBS (Gibco Lot #2557182). All transfections were performed 24 hours post plating by hepatocyte maintenance medium supplemented with 5% FBS (Gibco Lot #2557182) and LNPs containing gene editor mRNA and pegRNAs and/or ngRNAs targeting the relevant genomic locus. Amino acid sequences of gene editor proteins used in this study can be found in **Table IV**. LNPs were pre-equilibrated in media at 37°C for 5 minutes before dosing. After 48 hours of incubation, cells were lysed for genomic DNA extraction and editing analysis via next-generation sequencing (NGS) using QuickExtract DNA Extraction Solution (Biosearch Technologies #QE09050)

### Generation of Hep3b cell line bearing the p.R408W mutation

The Hep3B cell line harboring a PAH c.1222C>T mutation (Hep3B-R408W) was generated by EditCo Bio’s (formerly Synthego) proprietary CRISPR/Cas9 technology, single cell expansion and colony selection. Two homozygous Hep3B-R408W clones were identified and confirmed in house by Sanger sequencing and next-generation sequencing using the primers (fwd: 5’-CTTTCTCCAAATGGTGCCCTTCACT-3’ and rev: 5’-CGATGGTAGGGAAAGACAGTCTTCGATTA-3’). One representative clone A2 was expanded for use.

### Mouse model generation

The *Pah^R408W^*-UP model was obtained through a licensing agreement with the University of Pennsylvania, which provided two heterozygous male mice. The phenotypic characteristics of this model, including hyperphenylalaninemia and associated PKU-related traits, have been well documented in two prior publications [11,12].

This model was originally generated by the University of Pennsylvania in collaboration with Ozgene using homologous recombination in C57BL/6J embryonic stem cells. Specifically, exon 12 of the mouse Pah gene was replaced with the orthologous human exon 12, flanked by 500 base pairs of human genomic sequence upstream and downstream. The inserted human exon includes the c.1222C>T (R408W) mutation, which introduces an arginine-to-tryptophan substitution in the PAH enzyme on chromosome 10.

In parallel, an internal *Pah^R408W^* model was generated at Taconic Biosciences using CRISPR/Cas9 genome editing in C57BL/6NTac zygotes. This approach introduced the human R408W mutation into exon 12 of the endogenous Pah gene, mimicking the disease-associated allele while preserving murine regulatory architecture. Based on NCBI transcripts NM_008777.3 (mouse) and NM_000277.3 (human), this strategy enabled partial humanization of the gene by replacing 100 base pairs of mouse genomic sequence (50 bp upstream and 50 bp downstream of codon R408) with the corresponding human sequence. Exon 1, including the translation start site, remained unmodified. CRISPR reagents-including Cas9 protein, guide RNAs (gRNAs), and repair oligonucleotide-were microinjected into zygotes for germline editing.

To generate a non-disease control line, a second humanized mouse model (Pah^R408^) was engineered using the same CRISPR-based strategy, except without introducing the R408W mutation. These animals retain the humanized exon 12 sequence but encode the wild-type R408 residue. This model permits evaluation of human-mouse chimeric PAH expression in the absence of the pathogenic mutation and does not exhibit phenylalanine elevation or other PKU phenotypes.

Mice were monitored weekly for general health and signs of distress. Animals exhibiting significant discomfort, including >20% body weight loss, were euthanized by CO₂ asphyxiation in accordance with ethical guidelines. All animal procedures were conducted in compliance with Moderna Inc.’s Animal Care and Use protocols and approved by the Institutional Animal Care and Use Committee (IACUC).

### In Vivo LNP Administration and Sample Collection

Lipid nanoparticles (LNPs) were administered via lateral tail vein injection to 8-week-old female C57BL/6 mice (Jackson Laboratory) or to various phenylketonuria (PKU) mouse models (*Pah^R408W^*-UP, *Pah^R408W^*, Pah^R408^). Mice received LNPs at doses of 1 or 3 mg/kg, as described in the respective study designs. At designated time points, liver tissue (left lateral lobe) was harvested for DNA extraction (for indel quantification by next-generation sequencing, NGS) and protein analysis (Pah and Cas9 protein levels, quantified via Jess immunoassay). In parallel, plasma and whole blood were collected for phenylalanine (Phe) level measurements.

### Primary Mouse Hepatocyte isolation and culture

Primary mouse hepatocytes were isolated using the Miltenyi Biotec Liver Perfusion Kit. Briefly, fresh liver lobes (∼1.2 g) were perfused using the gentleMACS™ Octo Dissociator with Heaters, equipped with perfusion sleeves and perfusion-specific hardware. All reagents, including Enzymes D, R, and A, were reconstituted according to the manufacturer’s instructions and maintained as aliquots at −20°C to avoid freeze-thaw degradation. Perfusion involved sequential incubation with pre-digestion buffer, equilibration buffer, and an enzyme digestion mix. Liver tissue was secured onto the perfusion grid and subjected to a programmed perfusion cycle in the gentleMACS system at 37°C. After enzymatic digestion, liver tissue was transferred into a gentleMACS C-tube and dissociated. The resulting cell suspension was filtered through a 100 µm SmartStrainer and centrifuged at 30 × g for 5 minutes at 4°C to enrich hepatocytes. The cell pellet was resuspended in a warm plating medium [William’s E supplemented with 5% FBS, Dexamethasone, and Thawing/Plating Cocktail A (Gibco A15563)]. Cell viability and concentration were assessed via hemocytometer and ViaCell. Cells were seeded on collagen-coated plates with gentle rocking to ensure even distribution. After 4-6 hours of incubation, media was replaced with Maintenance Medium [William’s E + Dexamethasone + Cell Maintenance Cocktail B (Gibco A15564)] for subsequent culture.

### Cas9 Protein Quantification (Jess)

Frozen liver samples Hep3B cells were homogenized in RIPA buffer containing protease inhibitors. Protein concentration was normalized, and Cas9 expression was analyzed using the Jess automated western blot system (ProteinSimple) following manufacturer protocol with anti-Cas9 (Cell Signaling Tech. #19526S) antibodies. Vinculin (Cell Signaling Tech. #13901S) served as a loading control.

### Pah protein quantification (Jess)

Frozen liver samples were homogenized in RIPA buffer containing protease inhibitors. Protein concentration was normalized, and Pah expression was analyzed using the Jess automated western blot system (ProteinSimple) following manufacturer protocol with anti-Pah (Abcam. #ab178430) antibodies. GAPDH (Abcam. #ab8245) served as a loading control.

### Plasma sample collection, preparation, and phenylalanine analysis by LC-MS/MS

Whole blood samples (20∼30 µL per time point) were collected on Day −4 (baseline) and Day 7 (post-treatment). Samples were dispensed into pre-chilled tubes containing dipotassium ethylenediaminetetraacetic acid (K₂EDTA) and centrifuged at 4 °C for 10 minutes at 1,300 × g. The resulting plasma (10∼15 µL) was harvested and stored at −20 °C until analysis.

Frozen plasma samples were thawed on ice and diluted with deionized water (1:10). An aliquot of 50 µL diluted plasma was spiked with isotopically labeled L-phenylalanine (¹³C₉, ¹⁵N) (Sigma, 608017) as an internal standard (IS). Samples were extracted with 80% acetonitrile containing 0.1% formic acid to precipitate proteins. Supernatants were transferred to plate for LC-MS/MS analysis.

Phenylalanine quantification was performed using an Agilent 1200 HPLC system equipped with a Poroshell 120 HILIC column (1.9 µm, 2.1 × 150 mm; PN 693675-901, SN USJTR01019, LN S16023). The analyte and internal standard were detected using an Agilent 6470 triple quadrupole mass spectrometer operating in positive electrospray ionization (ESI+) mode.

### R-loop assays

Guide RNA solutions (IDT) were denatured in reaction buffer containing 0.1 mM MgCl2 (Fluka), 100 mM NaCl (Fluka), 50 mM Tris-HCl pH 8 (Fisher), and 100 μg/mg rBSA (New England Biolabs) by heating to 95°C for two minutes, then immediately chilling to 4°C. Complementary target DNA strands (Genscript) labeled with Cy5 or Cy3 were annealed at 10 μM in IDT duplex buffer by heating to 95°C for 2 minutes, then decreasing by 2°C/minute to 25°C, then holding at 4°C. DNA and guides were combined in reaction buffer to a concentration of 20 nM target DNA and 125 nM gRNA. 30 μL of the nucleic acid solution was added to a 384 well Corning solid back microplate, which was transferred into a BioTek plate reader. Cas9 nuclease (IDT) was diluted in reaction buffer to 100 nM,and 30 ul Cas9 solution was added to individual wells using a BioTEK autodispenser. After Cas9 addition to a well,the Cy3 (579 nm) and Cy5 (681 nm) emission in response to Cy3 excitation (540 nm) were measured at ∼1.7 second intervals while the solution was held at 25C. FRET was calculated as [Cy5 emission]/[Cy3 emission + Cy5 emission].

### Cleavage assays

Guide RNA and target DNAs were denatured and annealed, respectively, as described above, and combined in reaction buffer to a concentration of 80 nM gRNA + 20 nM target DNA. Cas9 was diluted to 100 nM Cas9 in reaction buffer and 8 μl Cas9 solution was added to 8 μl of target DNA + gRNA solution at 25°C. After the desired timepoint, the reaction was quenched by the addition of 16 μl 30 mM EDTA + 60 mg/ml Proteinase K (Thermo Scientific) and incubated for 10 minutes at 37°C, then 10 min at 95°C. 1 μl of reaction was then combined with 9 μL of Hi-Di Formamide (Applied Biosystems) and 0.5 μL Liz 120 Genescan standard (Thermo), then analyzed on an Agilent SeqStudio.

### Statistical Methods

All data were analyzed using GraphPad Prism v11.0. Results are reported as mean ± standard deviation (SD). For comparisons between two groups, unpaired two-tailed t-tests were used. For multiple group comparisons, one-way ANOVA with post-hoc multiple comparisons was applied. Significance thresholds were defined as p < 0.05 (*), p < 0.01 (**), p < 0.001 (***) and p < 0.0001 (****).

## Acknowledgments

We thank Maria Lei-Zhang, Ramita Kommuru and Yue Hui for their support throughout this study.

## SUPPLEMENTARY TEXT

### Generation and correction of other in vitro model cell lines for additional liver monogenic disease targets

We also generated Hep3B cell lines harboring additional common pathogenic mutations in the *PAH* gene, as well as mutations in *CBS* [31] and *G6PC* [32] that are implicated in inherited metabolic diseases (**Table II**). Mutations in *CBS* cause classical homocystinuria, a recessive disorder of sulfur amino acid metabolism characterized by elevated homocysteine levels and systemic complications including thromboembolism, developmental delay, and connective tissue abnormalities. Mutations in *G6PC* underlie glycogen storage disease type Ia (GSD-Ia), a severe pediatric disorder marked by fasting hypoglycemia, hepatomegaly, lactic acidosis, and failure to thrive.

We rationally designed 5 pegRNAs to template corrections of each mutation, in combination with various candidate ngRNAs. In **Supplementary Figure 2,** we present pegRNA screening results, with and without ngRNAs, for two clinically relevant PAH variants: R261Q (c.782G>A) and R243Q (c.728G>A). Several pegRNAs achieved promising levels of correction, with prime editing efficiencies reaching up to 20%. As shown in **Supplementary Figure 3**, we performed an initial screen of pegRNA and nicking guide **(Table III)** combinations for correction of the *CBS* I278T (c.833T>C) and *G6PC* Q347* (c.1039C>T) mutations. For *CBS*, editing efficiencies ranged up to ∼5% in the best-performing pegRNA/sgRNA pairs, demonstrating low but detectable activity at this locus. In contrast, screening for *G6PC* Q347* correction revealed several pegRNA combinations with markedly higher activity, including edits exceeding 30% efficiency in multiple conditions, highlighting this target as a promising candidate for therapeutic prime editing. These findings support the feasibility of extending LNP-delivered prime editing strategies beyond PAH to additional monogenic liver disorders.

### Screening of sequence and modification variants of PAH R408W-targeted pegRNAs

The sequence and chemistry of the pegRNA’s 3’ extension (PBS and RTT) plays a key role in activity. Chemical modification of the 3’ extension is likely required for sufficient resistance to exo- and endonucleases [33, 34]. Optimizing the hybridization energetics of the 3’ extension is also critical. The PBS must hybridize to the nicked NTS with sufficient stability to effectively prime reverse transcription [29], but a PBS+RTT that hybridizes too tightly to the complementary 5’ spacer produces an “autoinhibited”, low activity conformation [35]. Therefore, we examined a variety of approaches to enhance prime editing efficiency via engineering of the 3’ pegRNA extension. Additionally, we observed that incorporation of standard chemical modification patterns used in sgRNAs [22] resulted in significantly decreased prime editing efficiency with this pegRNA. We therefore also explore variations in the spacer and scaffold of the pegRNA in an attempt to overcome this challenge. These strategies are detailed below.

One approach to enhancing the 3’ stability of pegRNAs is to fuse an RNA-binding La protein to the prime editor, termed PE7, which helps protect the end of the pegRNA from degradation [36]. To utilize this approach, one must append a 3’ mU*mU*U sequence to the PBS, to enable La-binding. Surprisingly, upon doing so, we observed a significant decrease in prime editing activity compared to the previously used end-modified pegRNA (**Supplementary Figure 13**). We speculated that the addition of the terminal U’s was somehow altering the folding of the pegRNA and preventing binding to the prime editor protein.

Accordingly, we tested incorporating additional chemical modifications into this PBS designed to alter spacer:PBS binding, as well as pegRNA stability, and observed an increase in editing upon incorporation of 2’-fluoro modifications within the PBS (**Supplementary Figure 13)**.

We also speculated that, if incorporation of the terminal U’s increased spacer:PBS binding, that either shortening the PBS or lengthening the spacer might restore prime editing activity by influencing the folding of the pegRNA. However, in each case in which we tested shortened PBS sequences, we observed a reduction in prime editing efficiency (**Supplementary Figure 13)**. Similarly, upon testing pegRNAs bearing 1- and 2-nt longer spacers, we observed a decrease in prime editing efficiency (**Supplementary Figure 13)**. This is consistent with our findings showing reduced DNA cleavage and R-loop formation in *in vitro* assays with the corresponding sgRNAs (**Supplementary Figure 14**).

Additionally, although we observed minimal on-target undesired indels or scaffold incorporation following prime editing, we tested incorporation of a matched genomic sequence in the terminal hairpin of the pegRNA scaffold to prevent genomic mutations due to reverse transcriptase read through [9], identifying constructs which maintained prime editing efficiency while virtually eliminating genomic mutations due to scaffold incorporation (**Supplementary Figures 13 and 15**).

Lastly, we tested pegRNAs containing previously identified alternative nexus and/or linker nucleotide sequences within the guide scaffold. These sequences were previously identified in SELEX and shown to increase nuclease activity at some sites [37]. RNA folding analysis suggested the possibility of basepairing between the pegRNA 3’ extension and this region of the pegRNA scaffold, so we speculated that varying this region might enhance prime editing efficiency. Upon testing five such scaffolds, we were able to identify alternative sequences that enabled similar levels of prime editing to the original sequence, we failed to observe any increase in editing efficiency (**Supplementary Figure 5**).

## SUPPLEMENTARY FIGURES

**Supplementary Figure 1:**
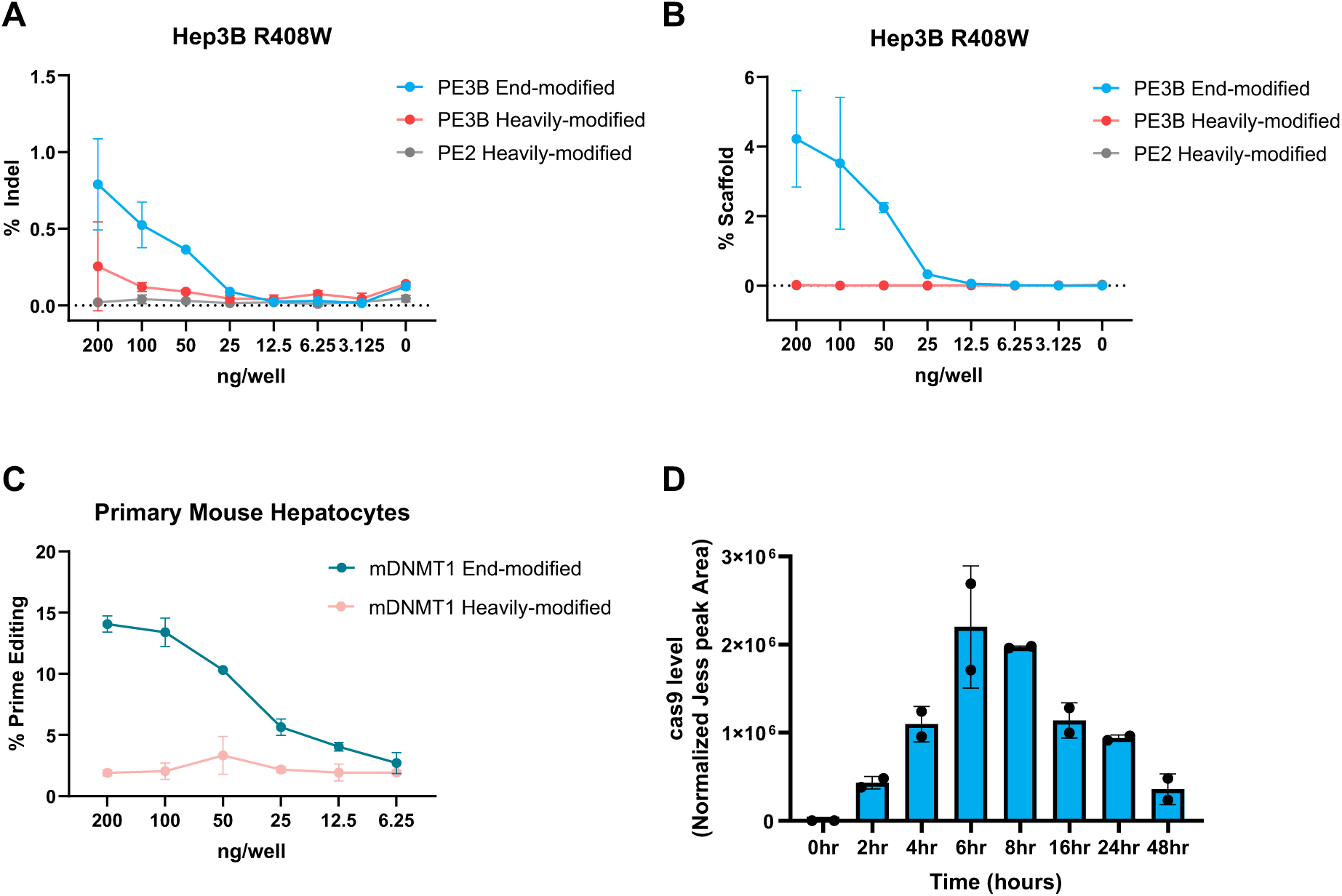
Dose-dependent editing, scaffold incorporation, and Cas9 expression in Hep3B R408W cells and Primary Mouse Hepatocytes. (A) Quantification of indel formation at the PAH locus in Hep3B cells harboring the homozygous R408W mutation following transfection with increasing doses of prime editor mRNA and gRNAs. Cells were treated with PE3b End-modified (blue circles), PE3b Heavily-modified (red circles), or PE2 Heavily-modified (gray circles) formulations using MessengerMax. Editing efficiency was quantified by amplicon sequencing at 72 hours post-transfection and reported as the percentage of indels. (B) Percent scaffold incorporation measured at the same target site and time point as in (A). (C) Prime editing efficiency at the mouse Dnmt1 locus in primary mouse hepatocytes treated with mRNA-LNP formulations. Cells were transfected with PE3b End-modified (blue circles) or Heavily modified (pink circles) pegRNA/sgRNA duplexes at decreasing total nucleic acid doses. Editing was quantified by amplicon sequencing 72 hours post-transfection. (D) Time course of Cas9 protein expression following LNP delivery of PE3b mRNA to Hep3B R408W cells. Protein levels were measured by Jess capillary Western blot at indicated time points and normalized to peak signal intensity. Cas9 expression peaked between 6-8 hours post-transfection and declined by 48 hours. Each bar represents the mean ± SD of biological replicates; individual data points are overlaid.

**Supplementary 2:**
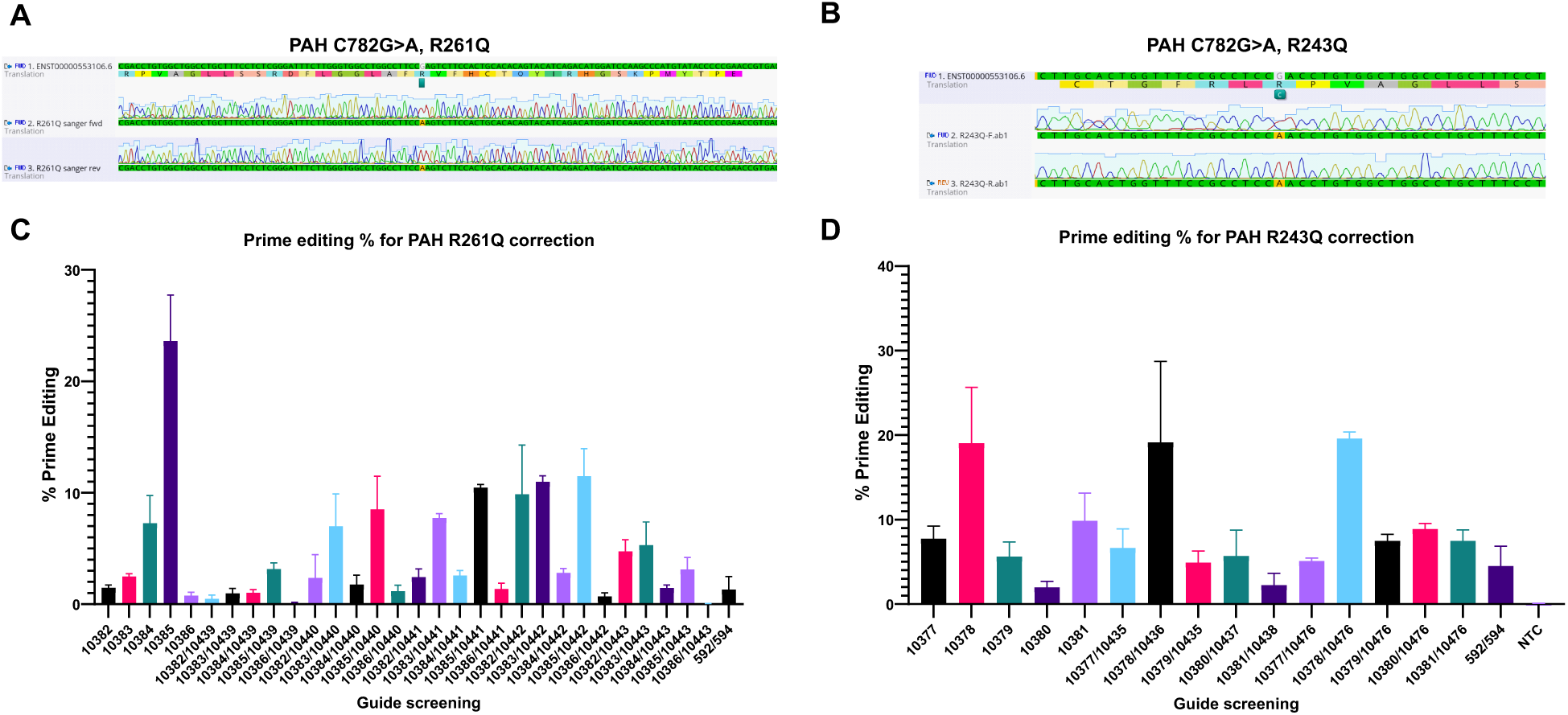
Sanger validation and pegRNA screening for prime editing correction of PAH R261Q and R243Q Mutations. (A-B) Sanger sequencing confirmation of installation of the homozygous PAH c.782G>A (p.R261Q) and c.728G>A (p.R243Q) mutations in engineered Hep3B cells. Forward and reverse sequencing traces for each variant confirm successful knock-in of the intended mutations at the endogenous PAH locus. (C) Screening of pegRNA candidates for correction of the R261Q variant. Hep3B-R261Q cells (Table VI) were transfected with a panel of pegRNAs and ngRNAs using the PE2 prime editor. Editing efficiency was assessed by amplicon deep sequencing at 72 hours post-transfection. (D) Screening of pegRNA candidates for correction of the R243Q variant. Hep3B-R243Q cells were transfected with the indicated pegRNAs and ngRNAs and mRNA encoding PE2. Editing efficiency was measured as in (C). Bars represent mean ± SD of prime editing efficiency from biological replicates.

**Supplementary Figure 3:**
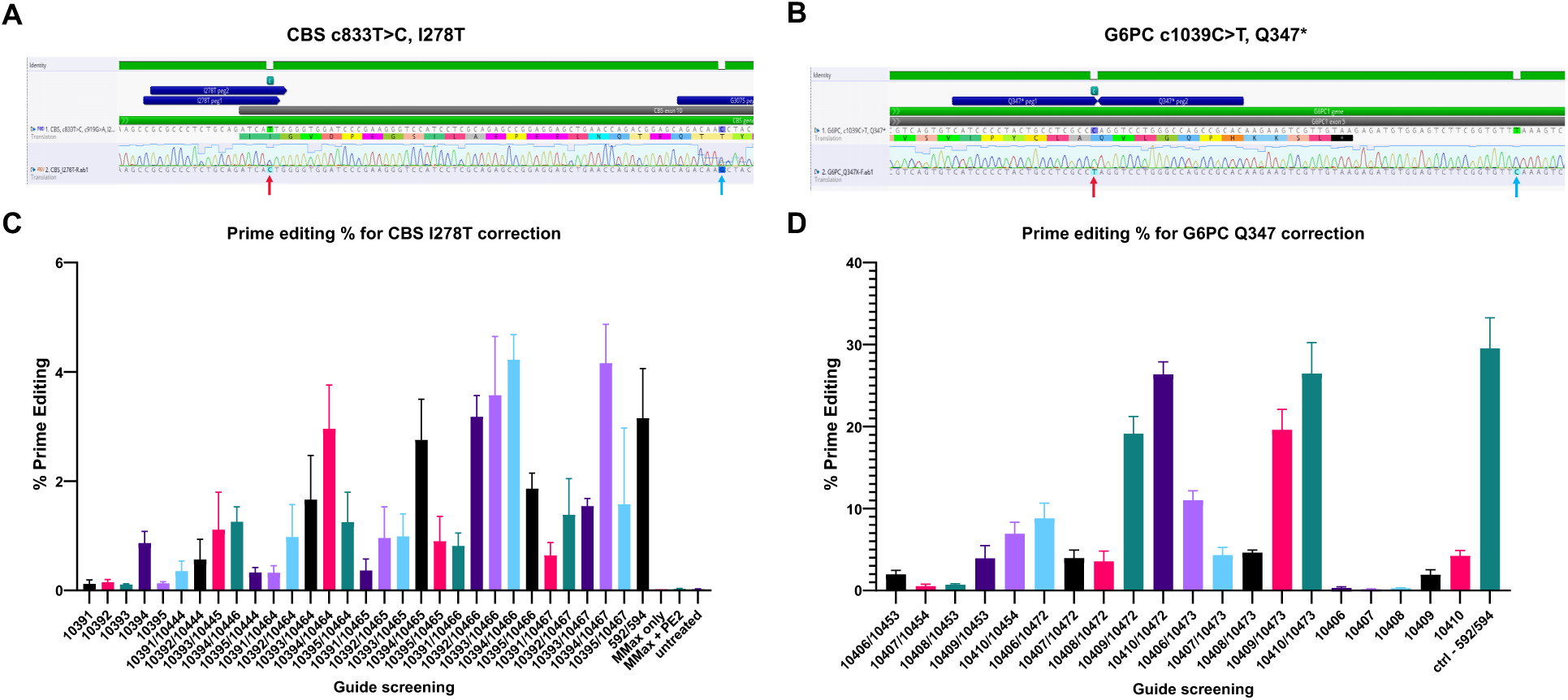
Sanger validation and pegRNA screening for prime editing correction of CBS I278T and G6PC Q347* mutations. (A-B) Sanger sequencing confirmation of installation of homozygous mutations introduced into Hep3B cells: c.833T>C (p.I278T) in the CBS gene (panel A) and c.1039C>T (p.Q347*) in the G6PC gene (panel B). Forward sequencing traces confirm accurate knock-in of the intended mutations. (C) pegRNA screening for correction of the CBS I278T variant (Table VI) and (D) G6PC Q347* variant. Engineered Hep3B cells were transfected with a panel of pegRNAs and nicking sgRNAs and mRNA-encoded PE2. Editing efficiency was quantified by amplicon sequencing 72 hours post-transfection. Bars represent mean prime editing efficiency across biological replicates. Several pegRNA/sgRNA pairs achieved editing efficiencies above 25%. Bars represent mean ± SD from biological replicates. Control = non-targeting control.

**Supplementary Figure 4:**
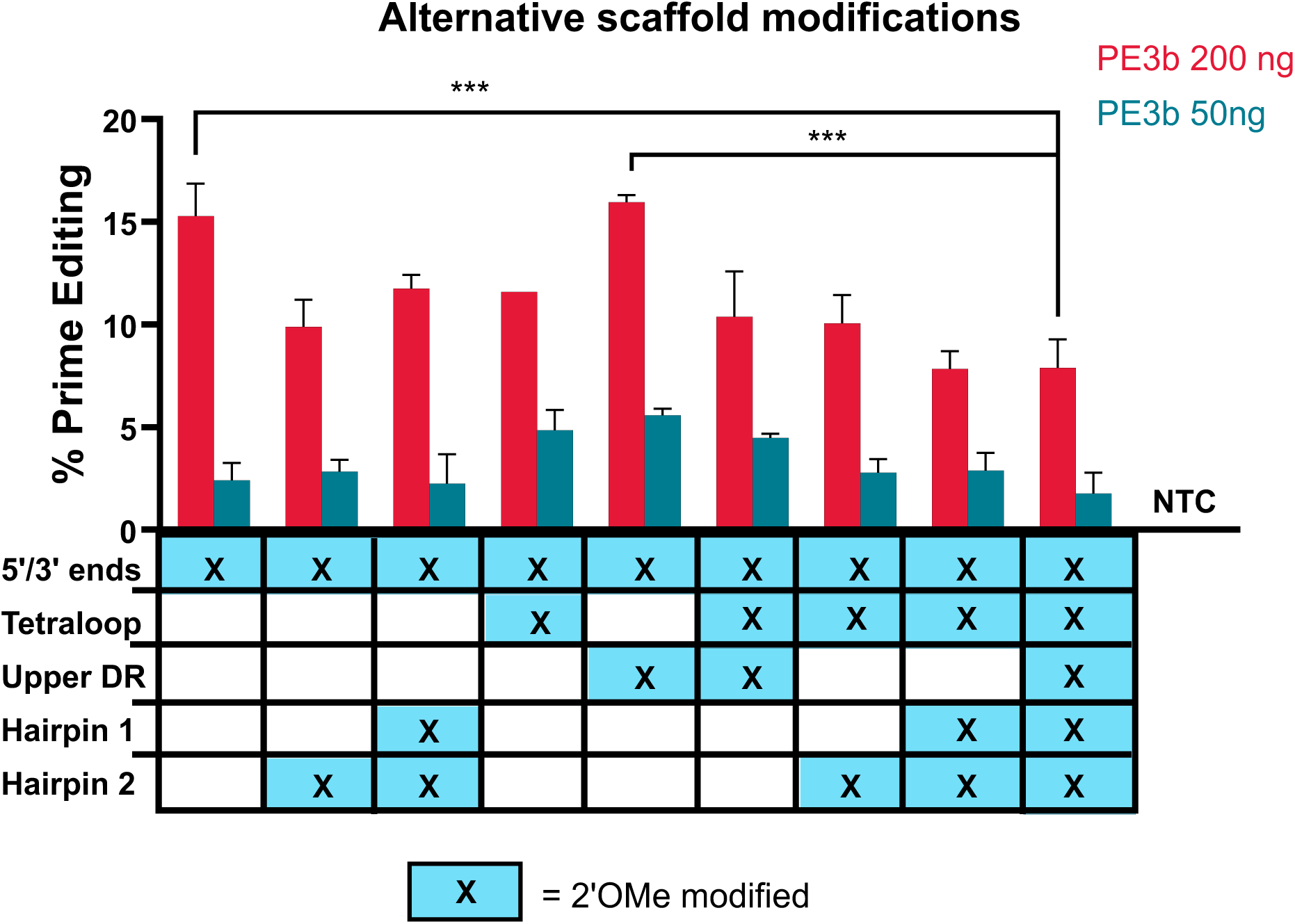
Prime editing with alternative scaffold modifications. We designed pegRNAs with a sequence identical to end-mod Pah R408W containing 2’OMe modifications at different combinations of the following site: the Tetraloop, the Upper Direct Repeat (DR), and Hairpins 1 and 2. The data above shows the PE3b editing efficiency of these guides when transfected into Primary Human Hepatocytes with Pah R408W-End mod-nicking guide, at 200 ng pegRNA/50 ng nicking guide (red) or 50 ng pegRNA/12.5 ng nicking guide (blue). Error bars represent standard error of two technical replicates. Ordinary one-way ANOVA with Dunnett’s multiple comparison test vs Pah R408W-heavily mod was performed. Significance levels: p < 0.05 (*), p < 0.01 (**), p < 0.001 (***). Blue filled boxes containing an “X” indicate the position of 2’OM2 modifications. Full sequences including modifications are provided in Table 1.

**Supplementary Figure 5:**
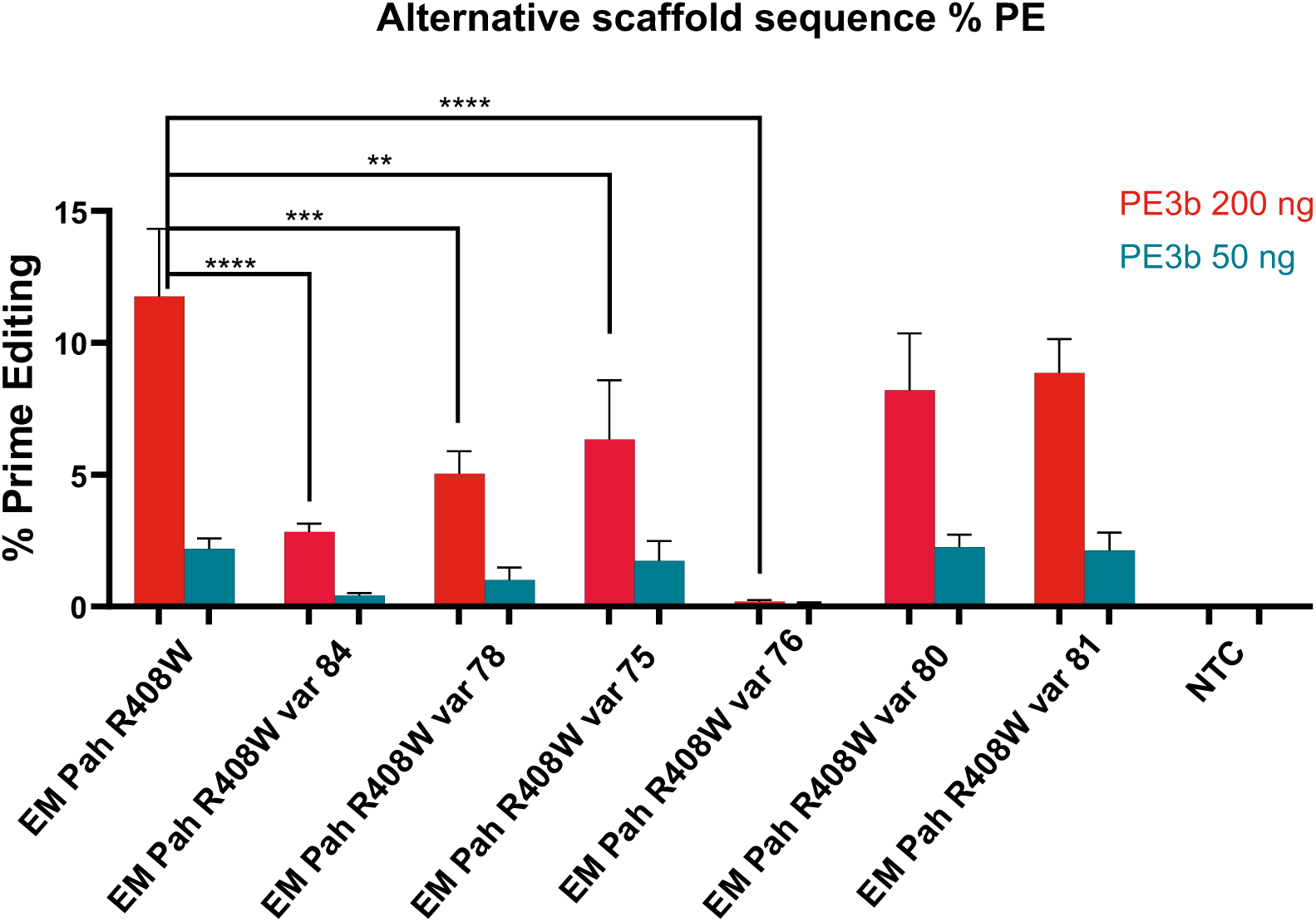
Prime editing with alternative scaffold sequences. We measured the prime editing activity of variants of end-modified PAH pegRNA containing alternative-sequence scaffolds previously demonstrated to improve cleavage activity [31] (Top) Alignments of the nexus and linker regions of these scaffold sequences showing positions of mutated sequences. Full sequences including modifications are provided Table 1. (Below)PE3b editing efficiency of these guides when transfected into Primary Human Hepatocytes with Pah R408W-End mod-nicking guide, at 200 ng pegRNA/50 ng nicking guide (red) or 50 ng pegRNA/12.5 ng nicking guide (blue). Error bars represent standard errors of three technical replicates. Ordinary one-way ANOVA with Dunnett’s multiple comparison test vs EM R408W was performed. Significance levels: p < 0.05 (*), p < 0.01 (**), p < 0.001 (***).

**Supplementary Figure 6:**
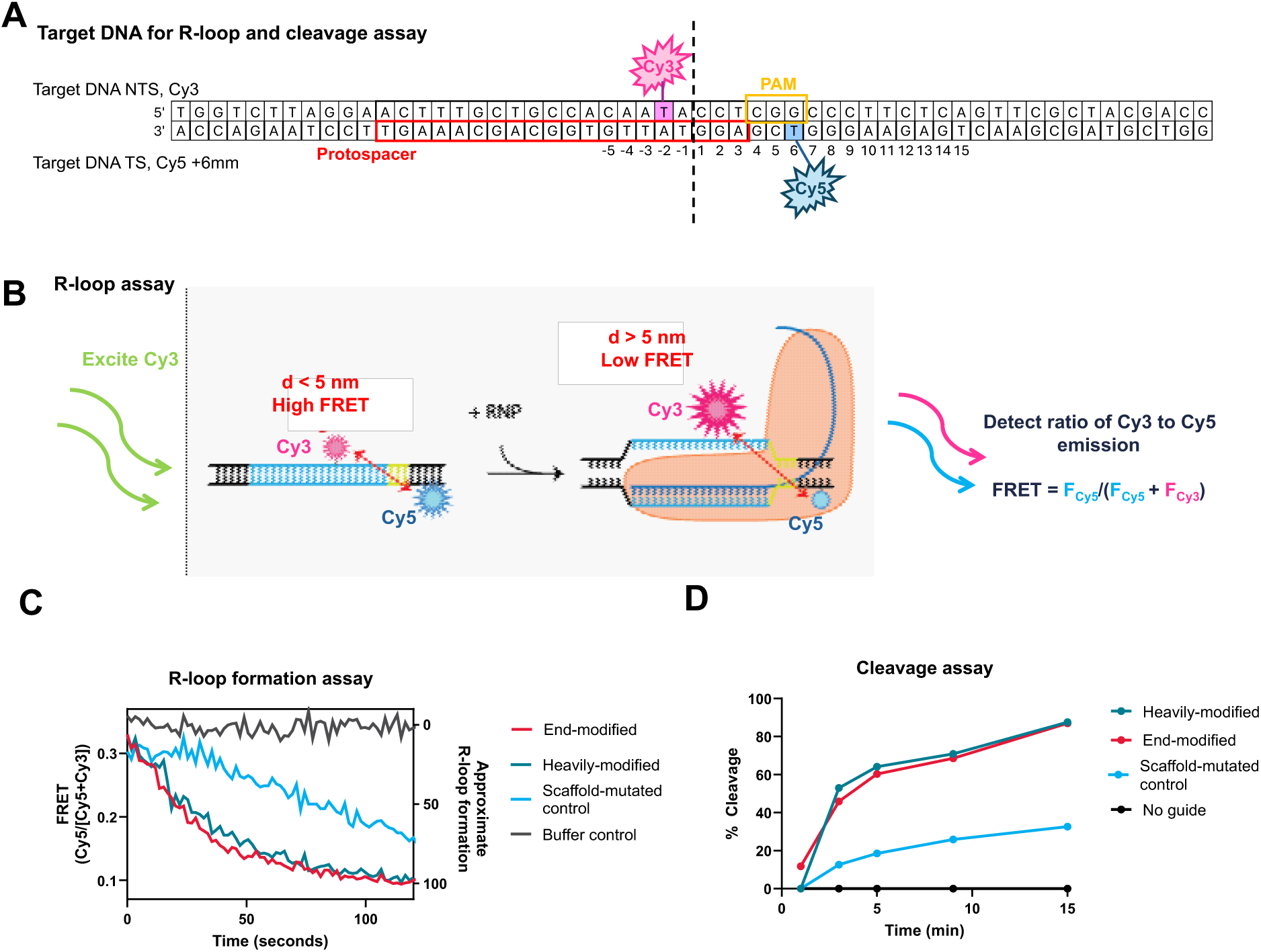
Biochemical assays for R-loop formation and DNA cleavage. (A) Cy3-Cy5 labeled target DNA used for PAH R408W biochemical experiments. Non-target strand is labeled with Cy3 opposite from the protospacer two nucleotides upstream from the cleavage site; and target strand is labeled with Cy5 six nucleotides downstream of the PAM. (B) Before R-loop formation, the NTS and TS are hybridized, and the Cy3 and Cy5 reporters are separated by <5 nm, producing high FRET and thus low Cy3 and high Cy5 emission in response to Cy3 excitation. R-loop formation displaces the non-target strand [24]. This increases the Cy3/Cy5 distance, decreasing FRET, and resulting in increased Cy3 and decreased Cy5 emission in response to Cy3 excitation. Cleaved DNA likewise produces low FRET, as the Cy3 and Cy5 are located in different cleavage products. (C) We measured R-loop formation over time after adding Cas9 to a solution of target DNA and sgRNA that was either (i) end-modified, (ii) heavily modified, or (iii) bearing a Cas9-binding impaired scaffold. FRET decreased over time as R-loop formation occurred. R-loop formation occurs with similar kinetics for end- and heavily-modified sgRNAs but is slower for binding-impaired sgRNAs. (D) Cleavage assay showing DNA cleavage after addition an RNP comtaining Cas9 and either of the three sgRNAs used in (C). As with R-loop formation, cleavage proceeds with similar kinetics for end- and heavily-modified sgRNAs but is slower for binding-impaired sgRNAs

**Supplementary Figure 7:**
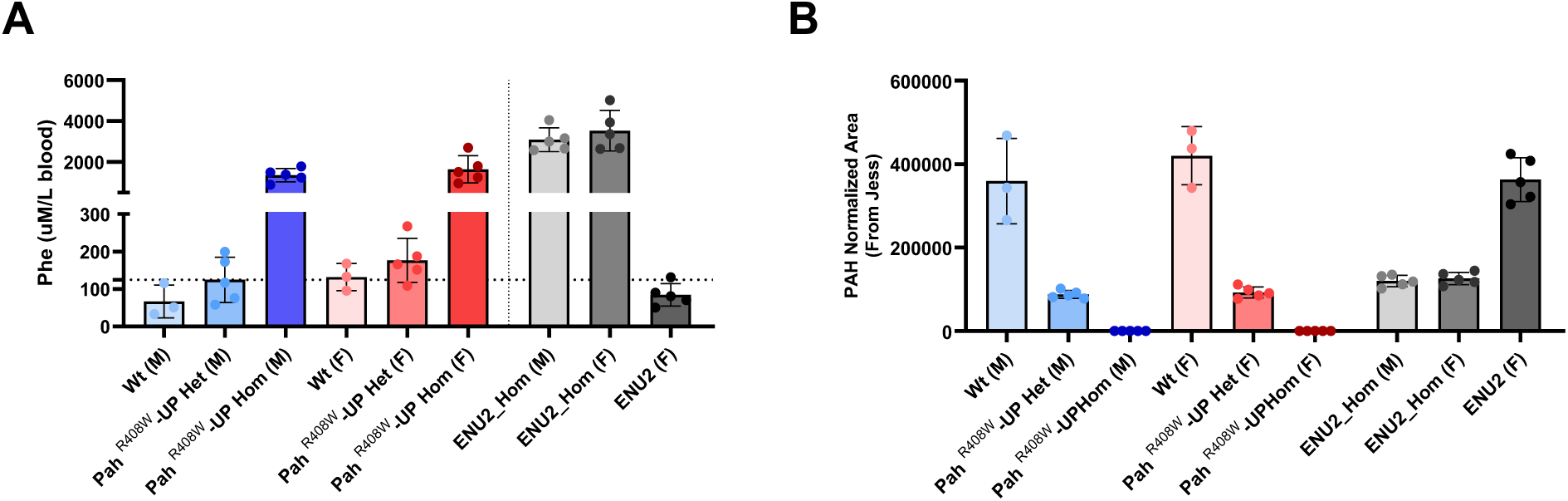
Characterization of Pah R408W mouse models from the University of Pennsylvania. (A) Blood phenylalanine (Phe) concentrations in wild-type (wt), heterozygous (Het), and homozygous (Hom) from the PAH ^R408W^-UP knock-in mouse models in comparison with male and female from the ENU2 line. Male (M) and female (F) animals are shown separately. Homozygous animals from both lines exhibited significantly elevated Phe levels compared to wild-type and heterozygous controls, exceeding 1200 µM in some cases. Dashed line indicates the upper threshold for therapeutic blood Phe concentration (∼360 µM). (B) PAH protein levels in liver tissue from the same animals, quantified by mass spectrometry and normalized to total proteome abundance. PAH expression was nearly undetectable in homozygous animals, while wild-type and heterozygous mice exhibited robust protein levels. Data represent mean ± SD from individual animals; each dot represents one biological replicate.

**Supplementary Figure 8:**
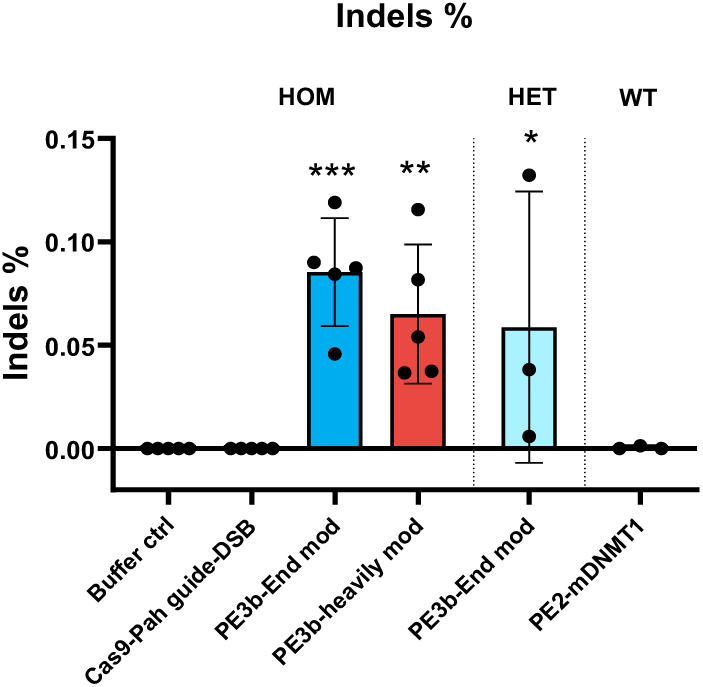
Minimal undesired on-target mutations were observed in Pah^R408W^-UP mice following in vivo Prime Editing treatment. Indel frequency at the Pah target locus in liver tissue from homozygous (HOM), heterozygous (HET), and wild-type (WT) Pah^R408W^-UP mice treated from Figure 2. Indel rates were quantified by deep amplicon sequencing of liver DNA one-week post-treatment and are reported as a percentage of total reads. All prime editing conditions resulted in low indel frequencies (<0.12%), and no scaffold-containing reads were detected across any treatment group (data not shown). Bars represent mean ± SD; each dot corresponds to an individual animal. Statistical significance was assessed by one-way ANOVA followed by Dunnett’s multiple comparisons test versus the buffer-treated control group. Significance levels: p < 0.05 (*), p < 0.01 (**), p < 0.001 (***).

**Supplementary Figure 9:**
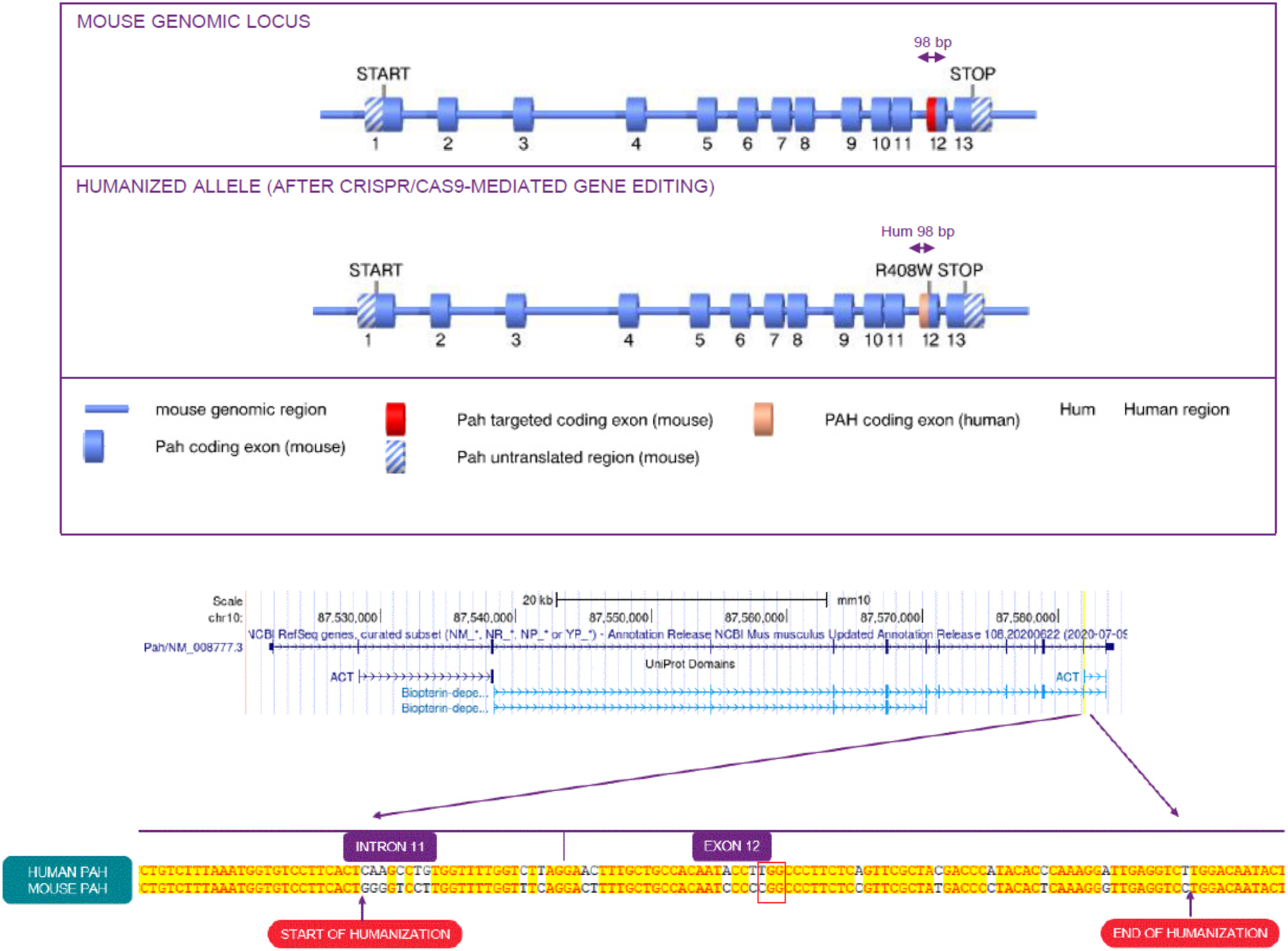
Design of the Humanized Pah^R408W^ knock-in mouse model. Top: Schematic of the mouse Pah locus (not drawn to scale) and the CRISPR/Cas9-edited allele. A 98-bp region spanning exon 12 and flanking intronic sequences was replaced with the corresponding human PAH sequence carrying the R408W mutation. Bottom: Sequence alignment showing the mouse-to-human transition in intron 11 and exon 12. The boundaries of the humanized region are highlighted. A control allele (Pah^R408^) lacking the R408W mutation was generated using the same approach.

**Supplementary Figure 10:**
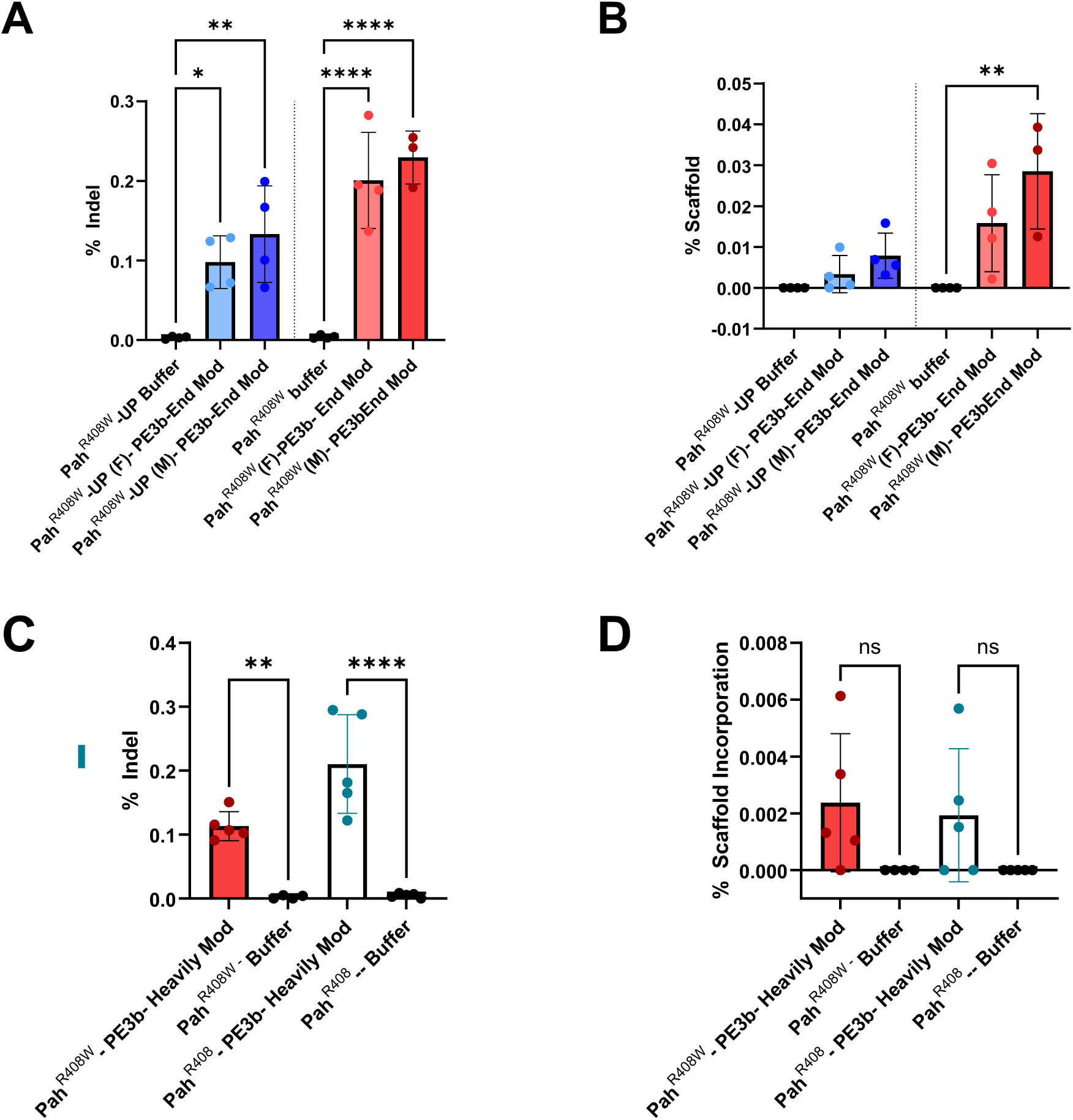
Indel and scaffold incorporation percentage in the editing comparison between the published in vivo model Pah^R408W^-UP and the internal knock-in transgenic model Pah^R408W^ and between the and control without point mutation Pah^R408^. (A-B) Comparison of undesired on-target editing events in Pah^R408W^-UP and Pah^R408W^ mice treated with LNP-delivered PE3b-End mod. (A) Indel frequencies and scaffold incorporation at the Pah locus in liver tissue were quantified by deep amplicon sequencing one-week post-treatment as described in Figure 3. (C-D) Comparison of undesired on-target editing events in Pah^R408W^ mice and in the control animals without the point mutation Pah^R408^ treated with LNP-delivered PE3b-Heavily mod guides. Data are presented as mean ± SD. Each dot represents one animal. Statistical significance was assessed using one-way ANOVA followed by Dunnett’s multiple comparisons test versus buffer-treated control. Significance levels: p < 0.05 (*), p < 0.01 (**), p < 0.0001 (****), ns = not significant.

**Supplementary Figure 11:**
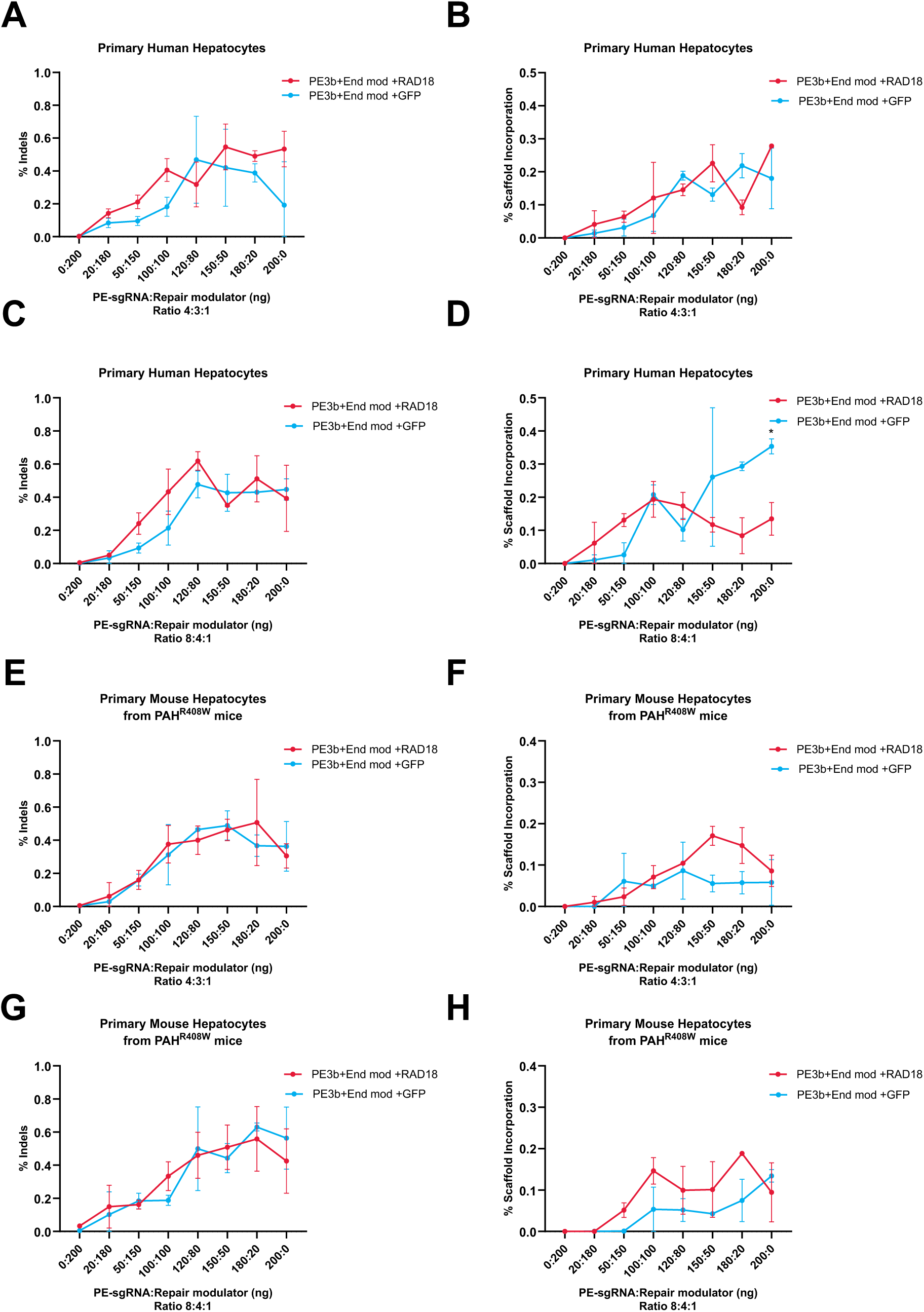
RAD18 enhances prime editing efficiency without increasing indel and scaffold incorporation in primary human and mouse hepatocytes. (A, C, E, G) % Indel formation and (B, D, F, H) % scaffold incorporation were measured following prime editing in primary human hepatocytes (A-D) and primary mouse hepatocytes derived from Pah^R408W^ mice (E-H). Experiments are as described in Figure 5. Cells were transfected with PE3b + end-modified sgRNA constructs and either RAD18 or GFP control mRNA, across a range of total RNA input amounts (up to 200 ng). Two RNA ratios of PE3b:sgRNA:repair modulator were tested: 4:3:1 (A-B, E-F) and 8:4:1 (C-D, G-H). Across multiple doses and in both human and mouse hepatocytes, RAD18 co-delivery does not lead to a notable increase in indel formation (A, C, E, G) and scaffold incorporation (B, D, F, H) compared to GFP controls. Data represent mean ± SD of technical replicates. Statistical significance was assessed by two-way ANOVA, followed by Dunnett’s or Sidak’s multiple comparisons test, as appropriate, comparing each concentration point to the GFP control group. Significance levels: p < 0.05 (*).

**Supplementary Figure 12:**
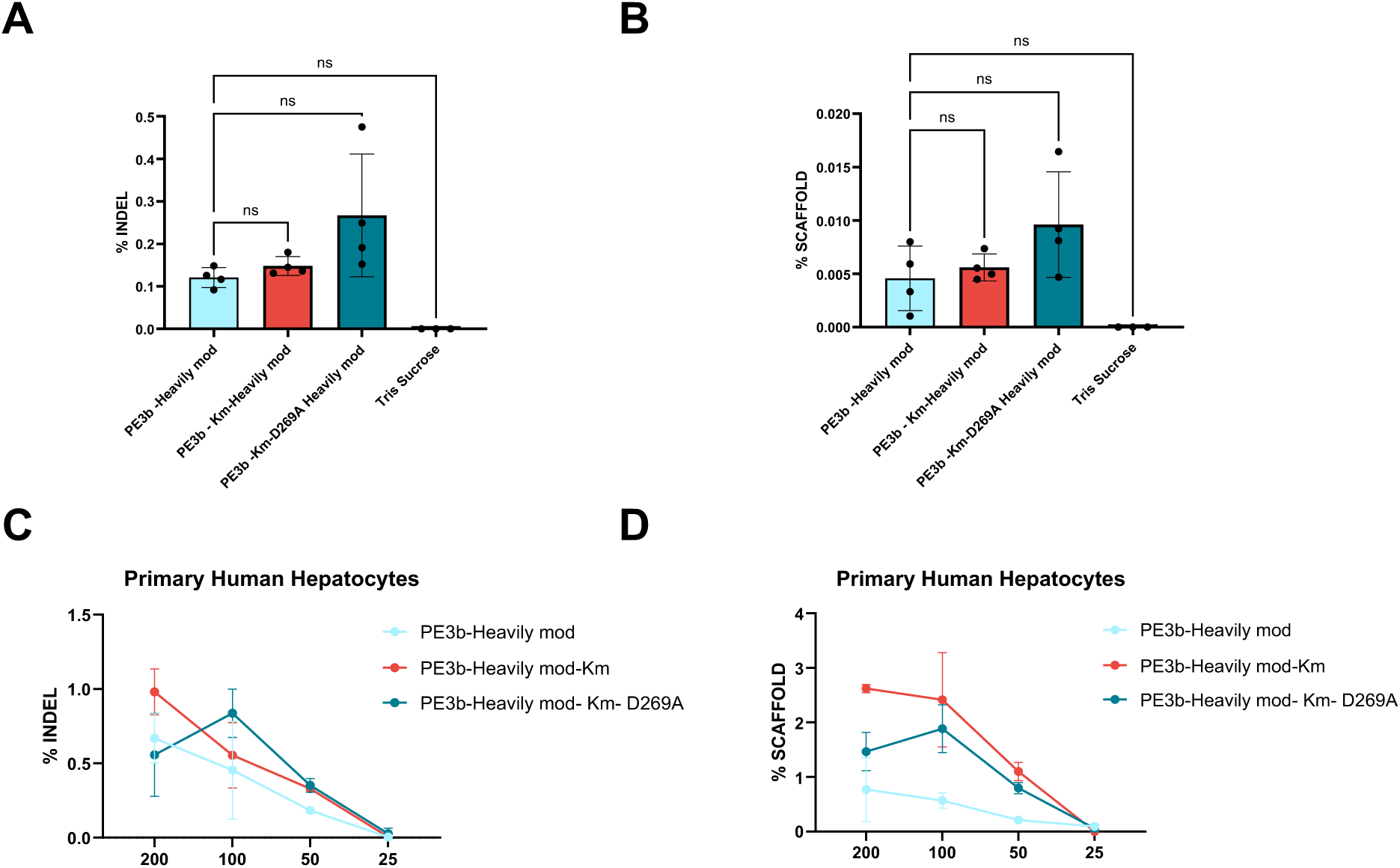
Impact of prime editor protein engineering on indel formation and scaffold incorporation. **(A-B)** In vivo prime editing was evaluated in heterozygous Pah R408W knock-in mice treated with a single intravenous dose (3 mg/kg) of LNP-formulated PE3b mRNA and chemically modified pegRNA/nicking gRNA. Mice were sacrificed one-week post-treatment as described in Figure 6. (A) Indel frequency at the PAH R408W locus was measured 72 hours post-transfection and expressed as percent of total reads. (B) Scaffold incorporation was assessed by quantifying pegRNA-derived scaffold sequences at the target site. (C-D) Dose titration study in PHH comparing indel formation and scaffold incorporation across PE3b variants. Cells were transfected with decreasing doses (200 ng to 25 ng) of total nucleic acid (editor mRNA + guides). (C) Percent indels and (D) percent scaffold reads were quantified by amplicon sequencing. Data are presented as mean ± SD. Each point represents a biological replicate. Statistical significance was assessed by one-way ANOVA followed by Tukey’s multiple comparisons test with ns = not significant.

**Supplementary Figure 13:**
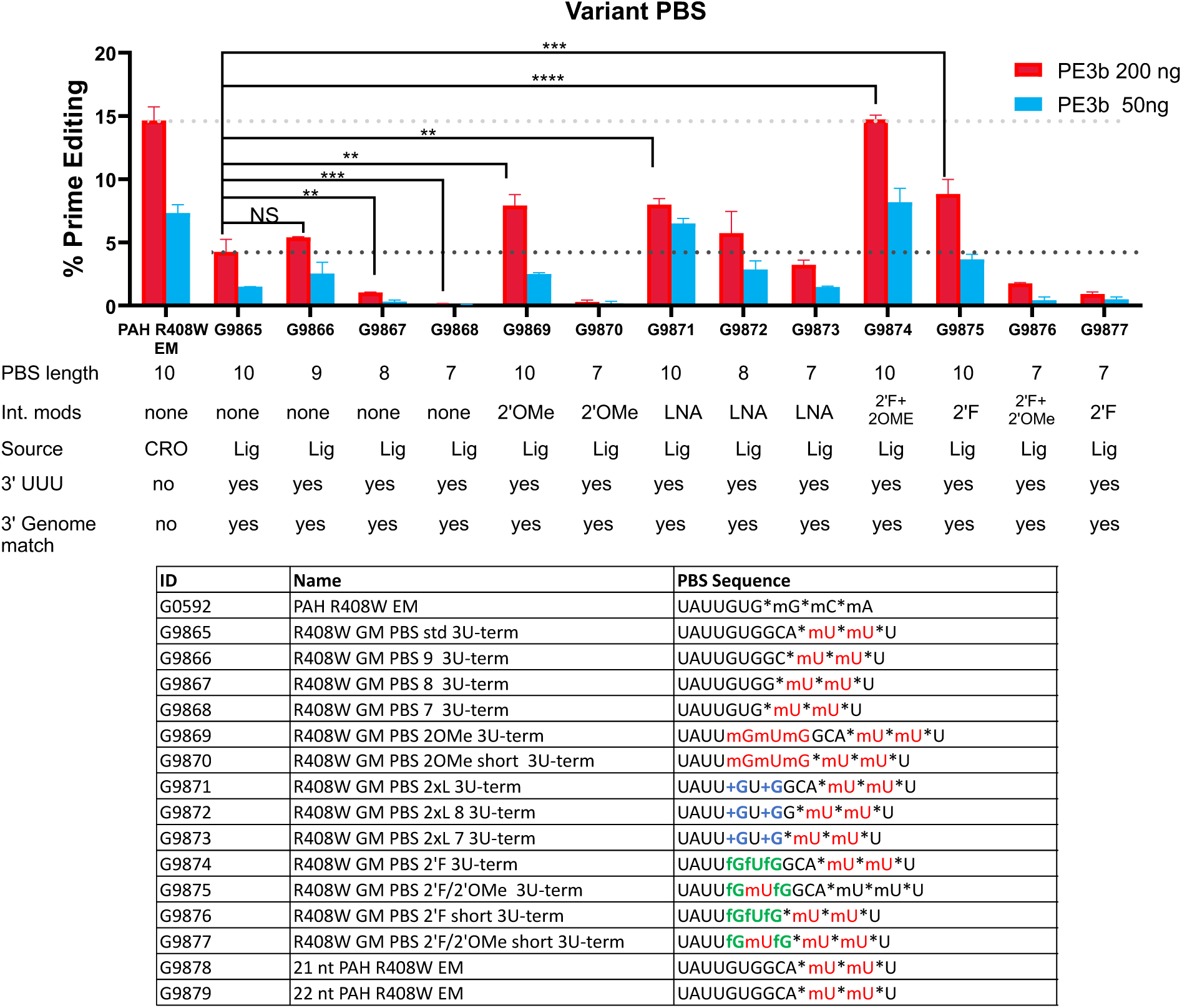
PE3b activity with variant PBSs. We designed pegRNAs otherwise identical to PAH R408W-End mod (G0592) but containing a genome match scaffold, a terminal 3’ mU*mU*U, and either no further modifications or a shortened PBS and/or 2’OMe, 2’Fluoro, 2’OMe + 2’Fluoro, or locked nucleic acid modifications in the PBS (Top) Data showing the PE3b editing efficiency of these guides when transfected into Primary Human Hepatocytes with Pah R408W-End mod-nicking guide, using either 200 ng pegRNA/50 ng nicking guide (red) or 50 ng pegRNA/12.5 ng nicking guide (blue). Error bars represent the standard error of two technical replicates. Ordinary one-way ANOVA with Dunnett’s multiple comparison test was performed. (Bottom) Table showing PBS sequences of the guides. “*” indicates a phosphorophioate linkage, “m” indicates a 2’-OMe modification, “f” indicates a 2’ fluoro modification, and “+” a locked nucleic acid. Full sequences including modifications are provided in Table 1. Significance levels: p < 0.05 (*), p < 0.01 (**), p < 0.0001 (****), NS = not significant.

**Supplementary figure 14:**
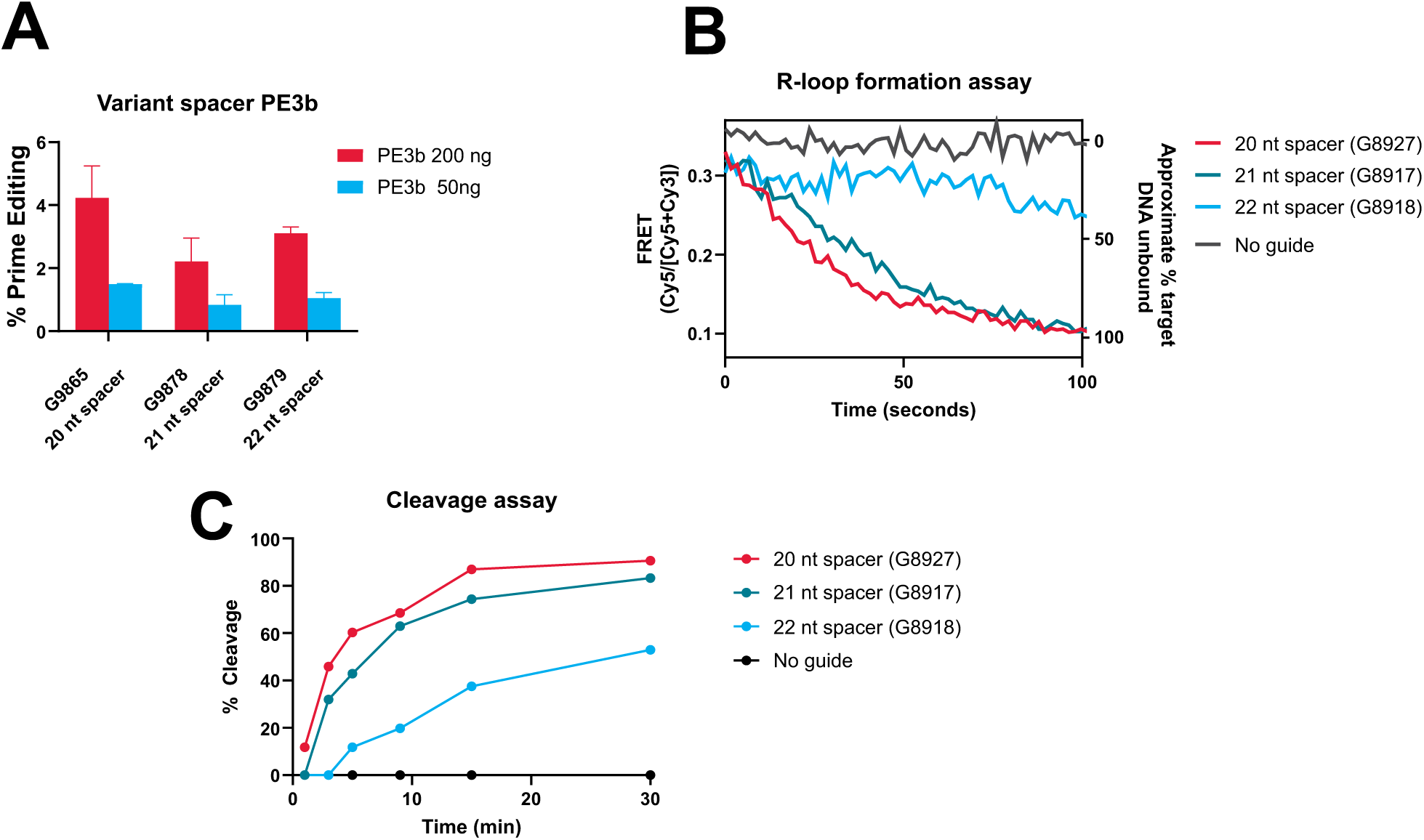
Prime editing and biochemical assay with extended spacers. (A) Data showing the PE3b editing efficiency of pegRNAs containing 20 nt, 21 nt, and 22 nt spacers when transfected into Primary Human Hepatocytes with Pah R408W-End mod-nicking guide, at 200 ng pegRNA/50 ng nicking guide (red) or 50 ng pegRNA/12.5 ng nicking guide (blue). Error bars are standard errors of two technical replicates. Ordinary one-way ANOVA with Dunnett’s multiple comparison test was performed, but for no samples did p <0.05. (B) R-loop formation assay and (C) kinetic cleavage assay with sgRNAs containing 20, 21, and 22 nt spacers.

**Supplementary Figure 15:**
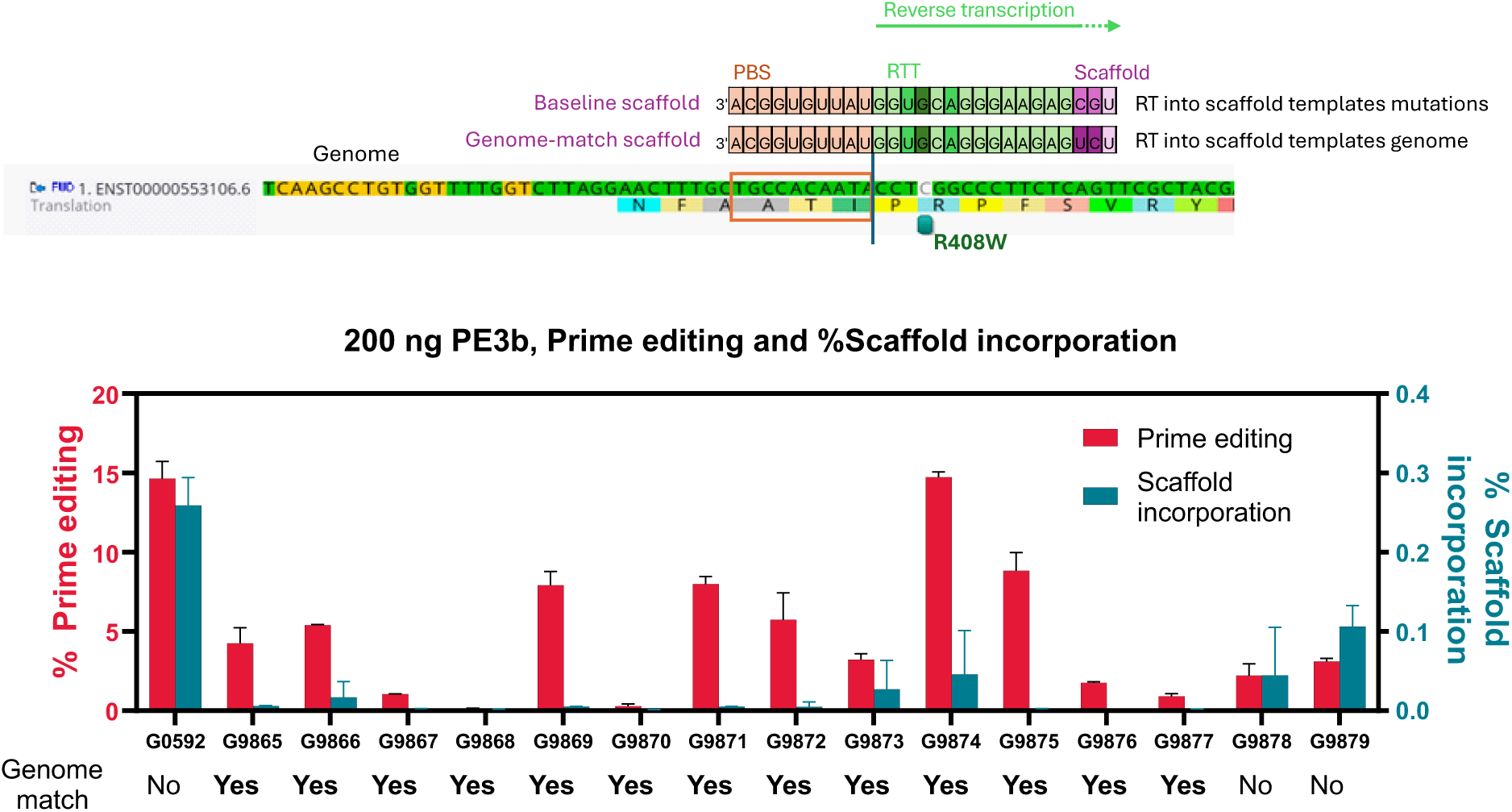
Genome match pegRNAs show decreased scaffold incorporation. (Top) Schematic showing baseline and genome match scaffolds. Reverse transcription into the baseline scaffold will template point mutations while reverse transcription up to three nucleotides into the genome match scaffold will template a perfect match to the genome and thus no additional mutations. (Bottom) Prime editing (red, left Y axis) and scaffold incorporation (blue, right Y axis) of pegRNAs with and without the genome match scaffold. Error bars represent two technical replicates.

## Supplementary Tables

**Table I.**
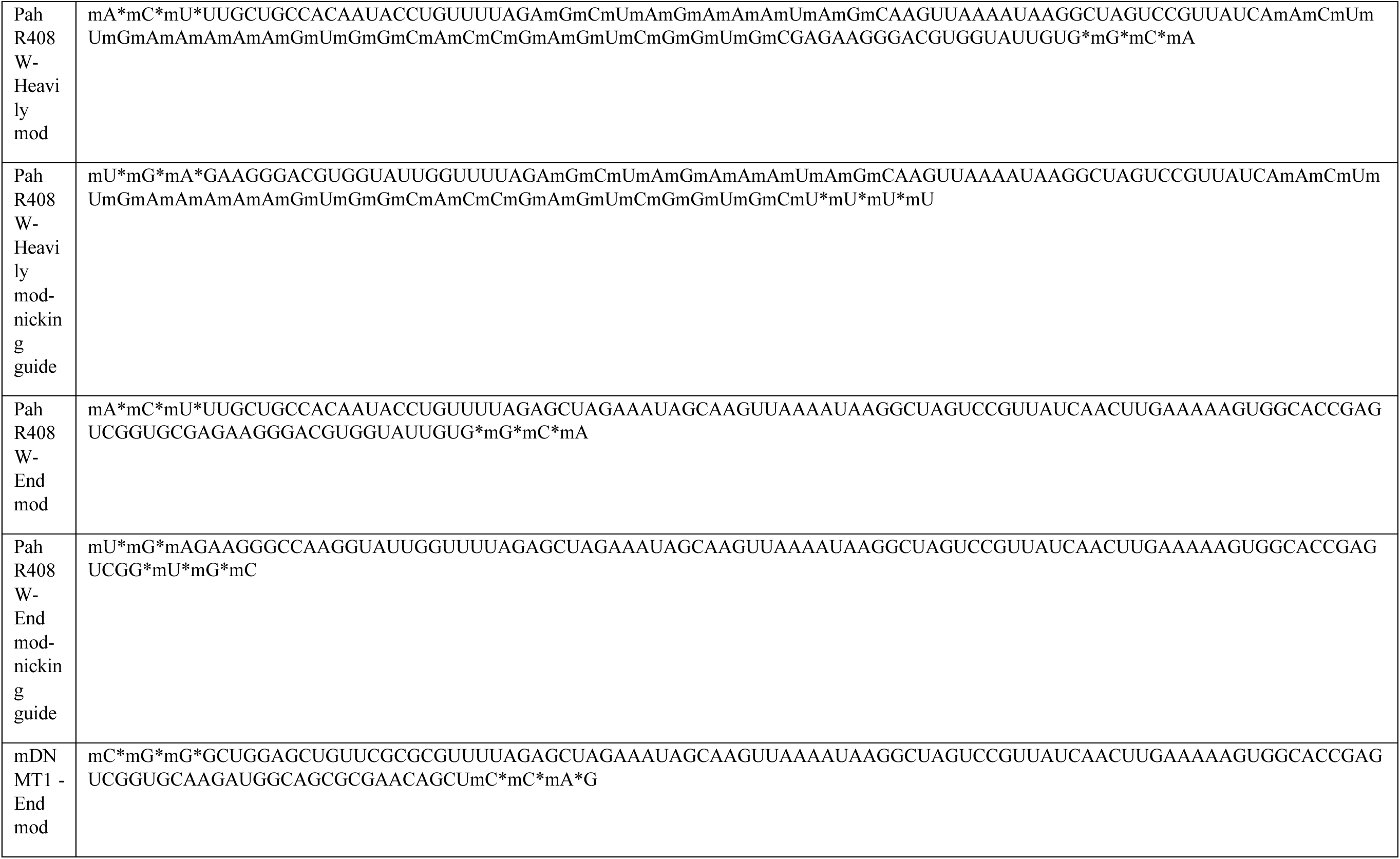
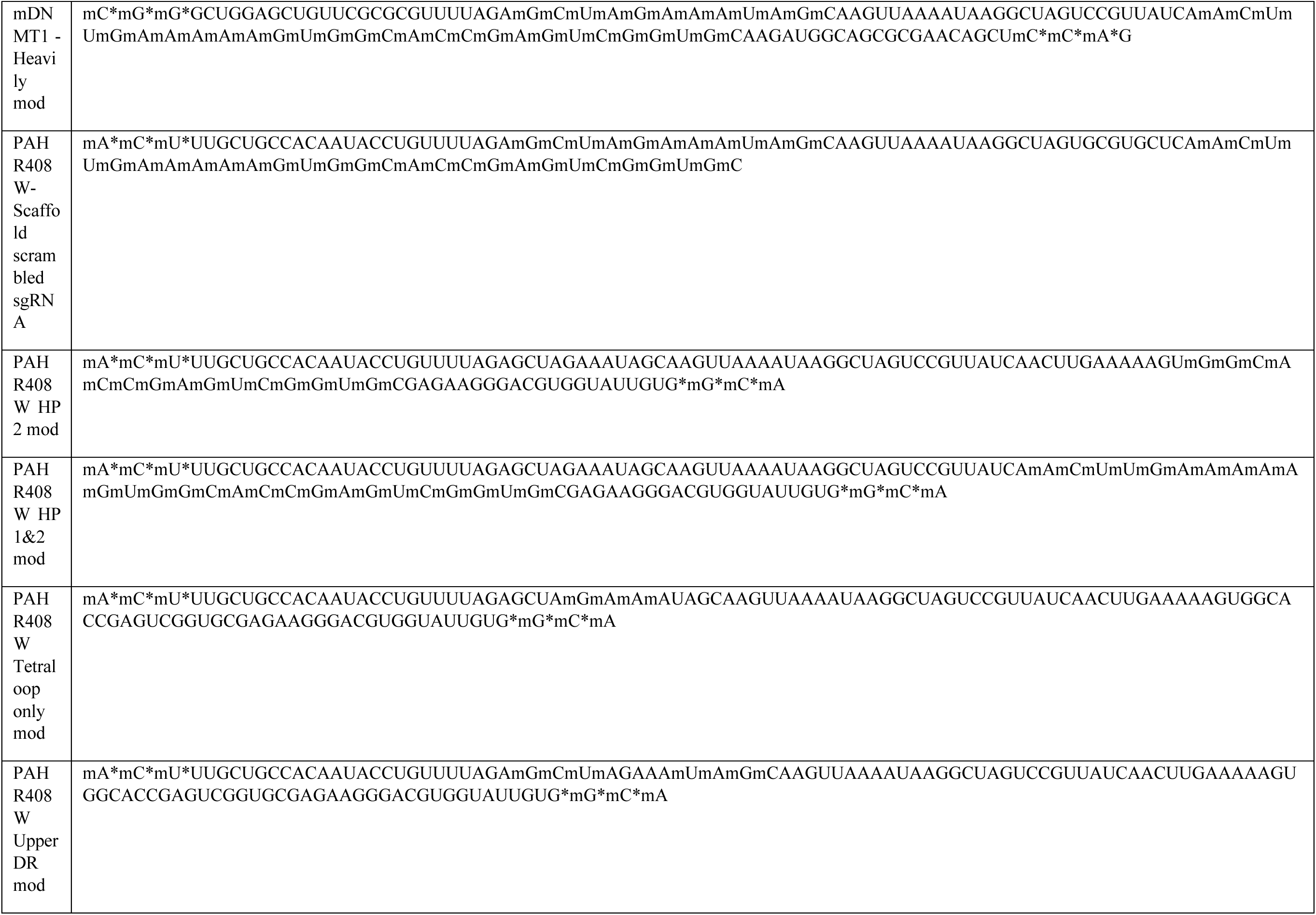
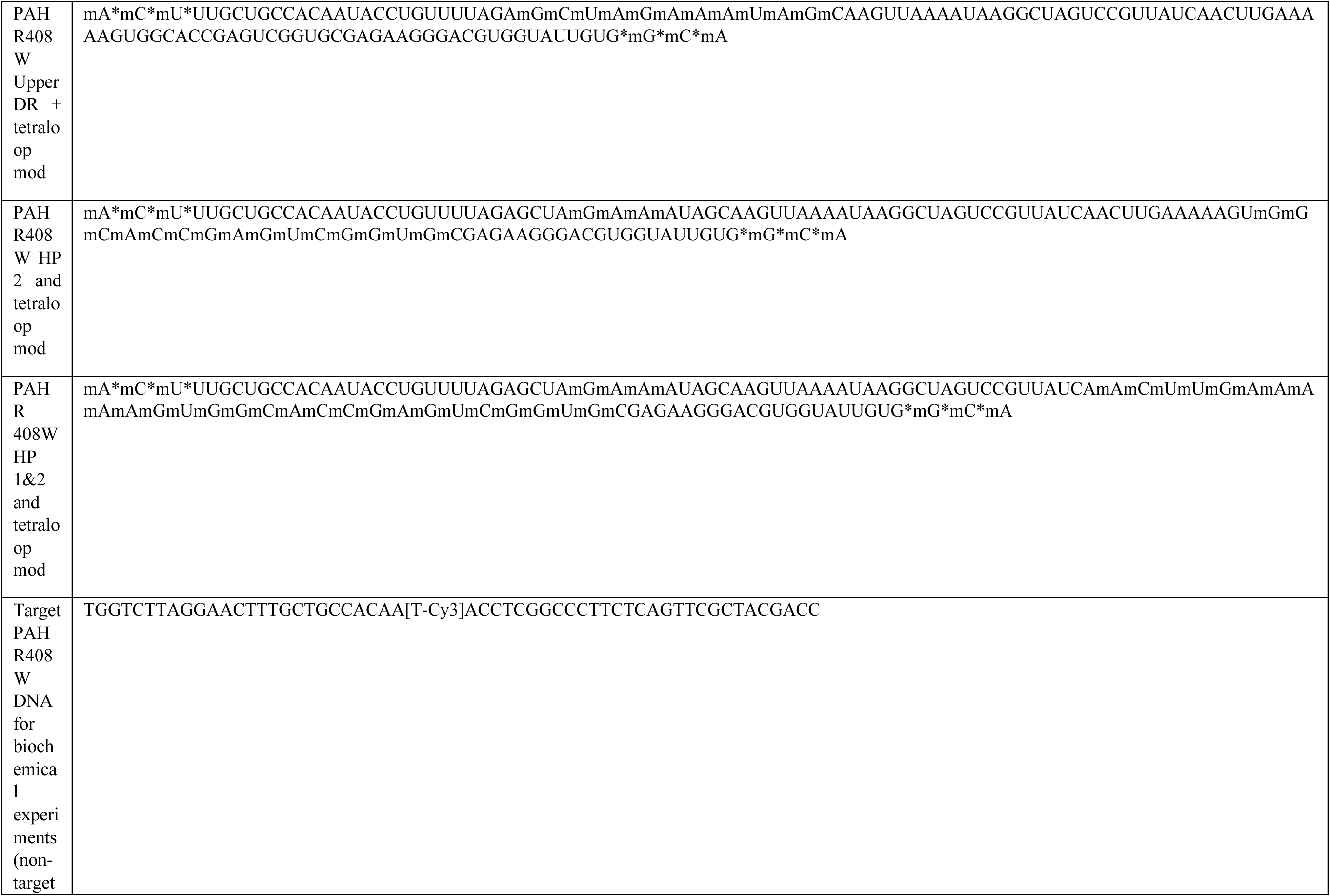
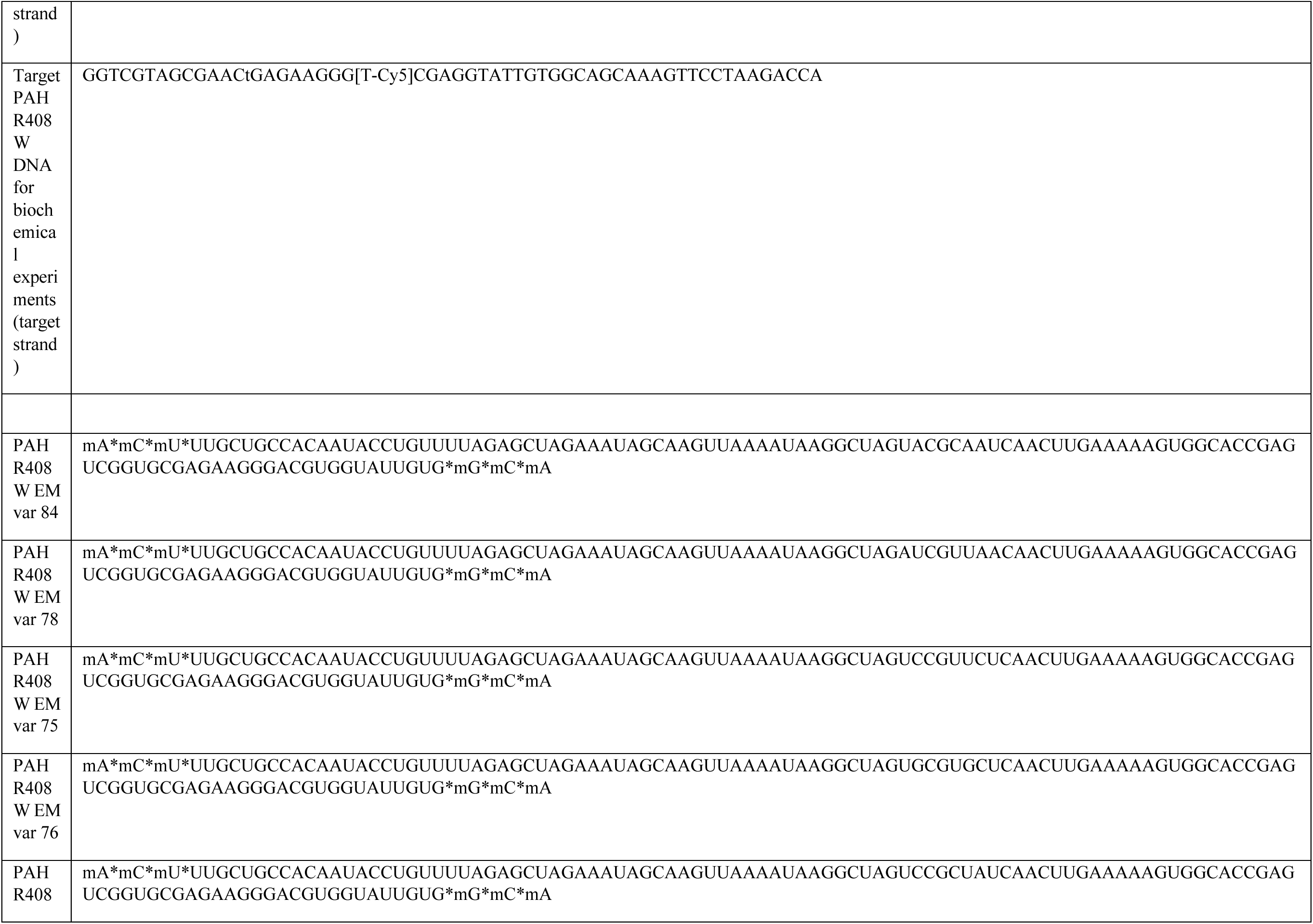
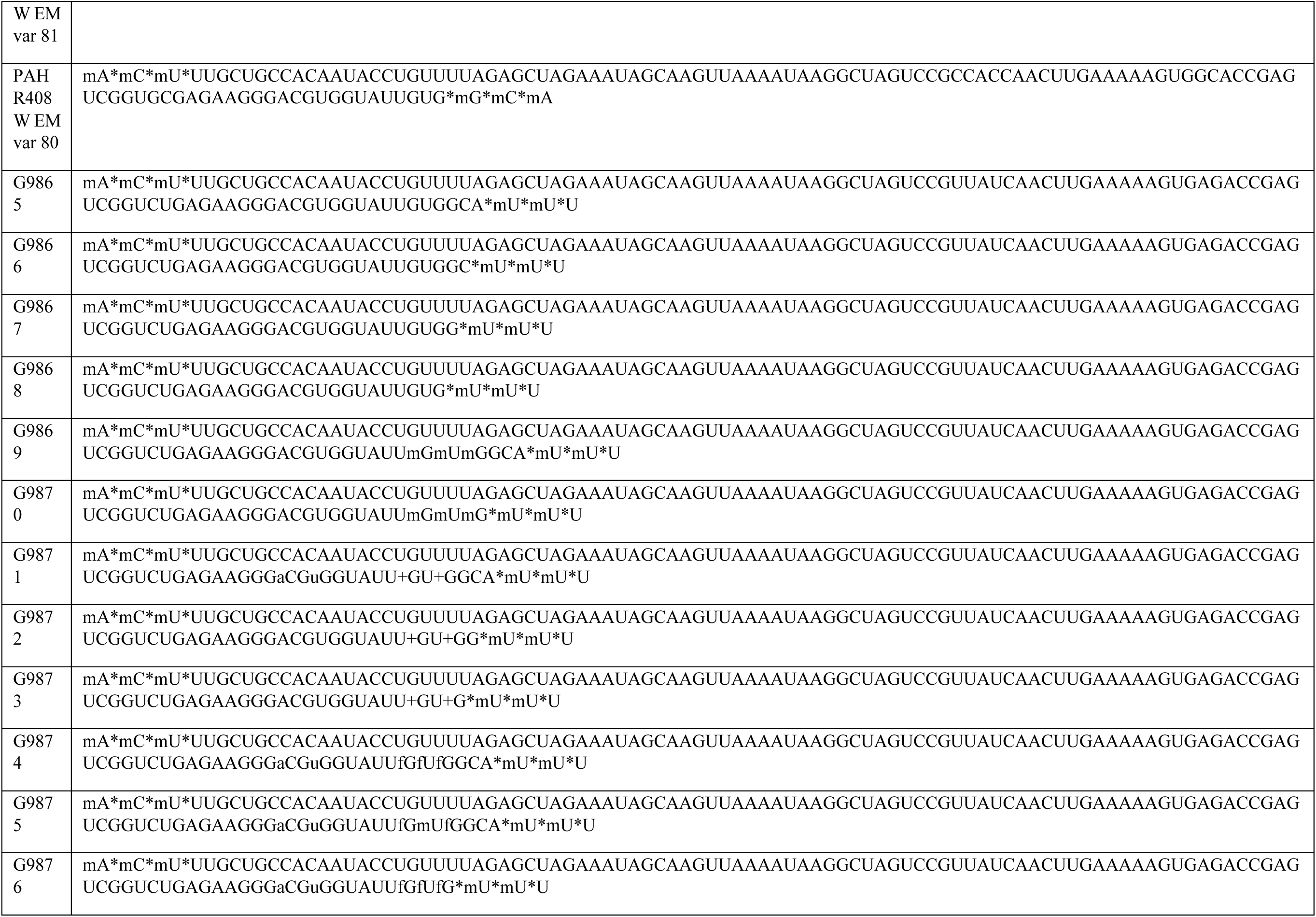
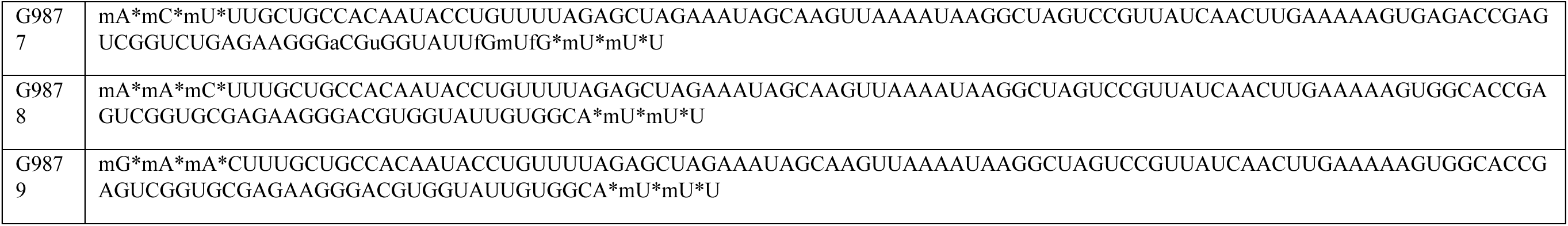
Sequences of sgRNAs and pegRNAs targeting PAH p.R408W.

**Table II.**
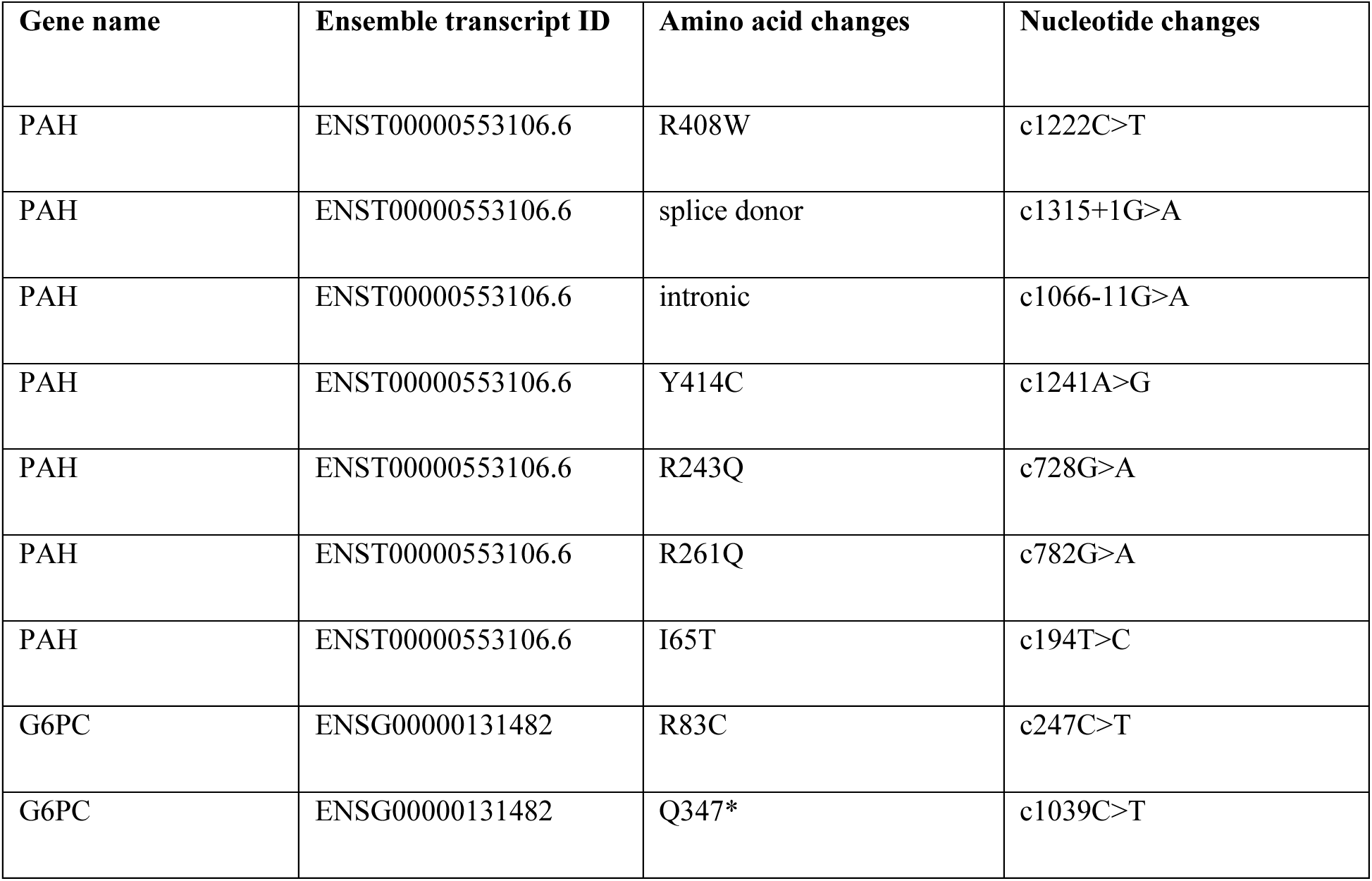
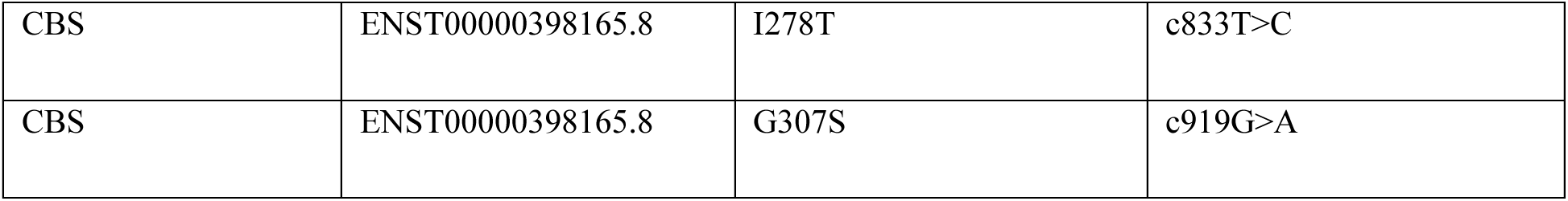
Mutations targeted for correction in this study.

**Table III.**
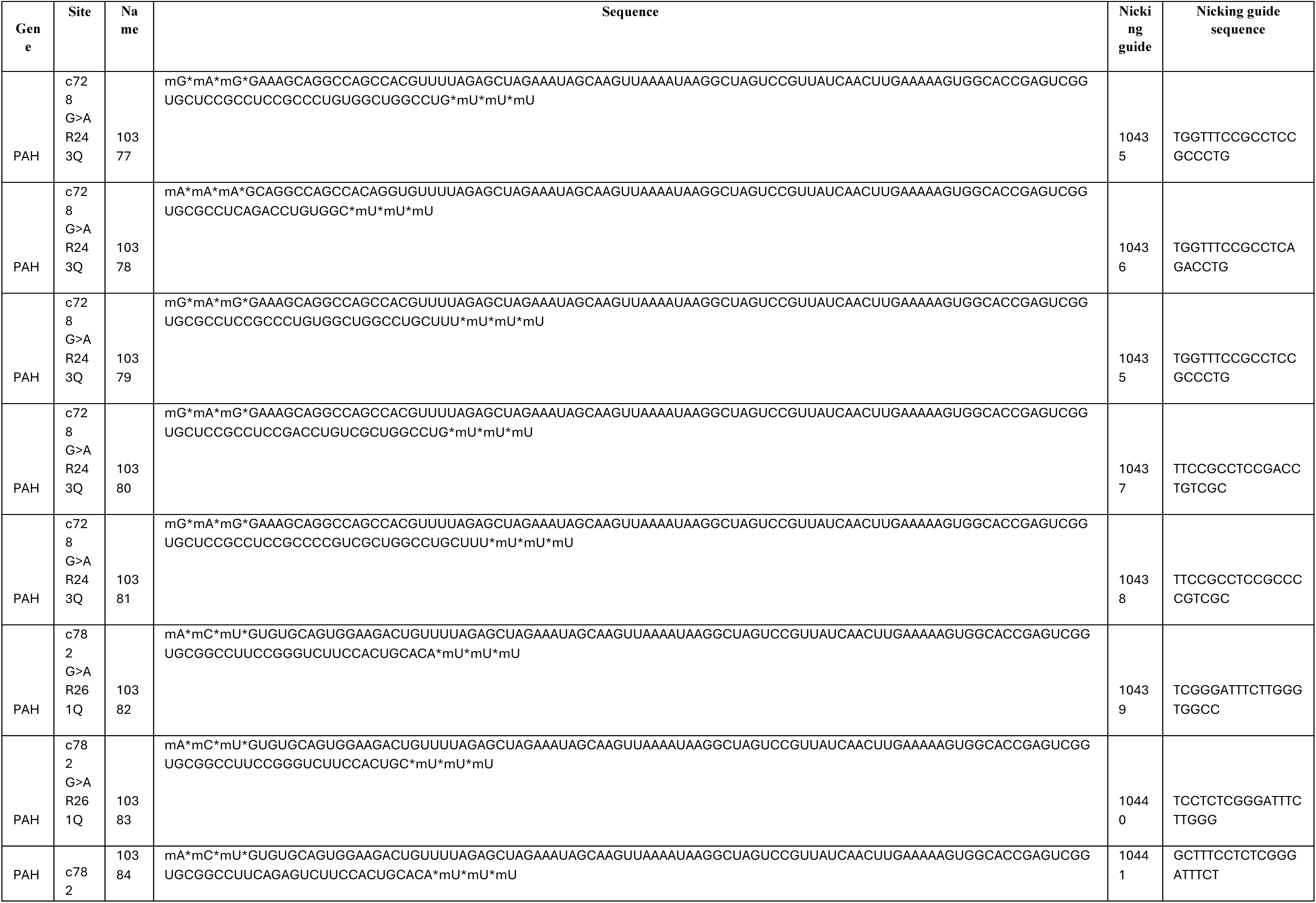
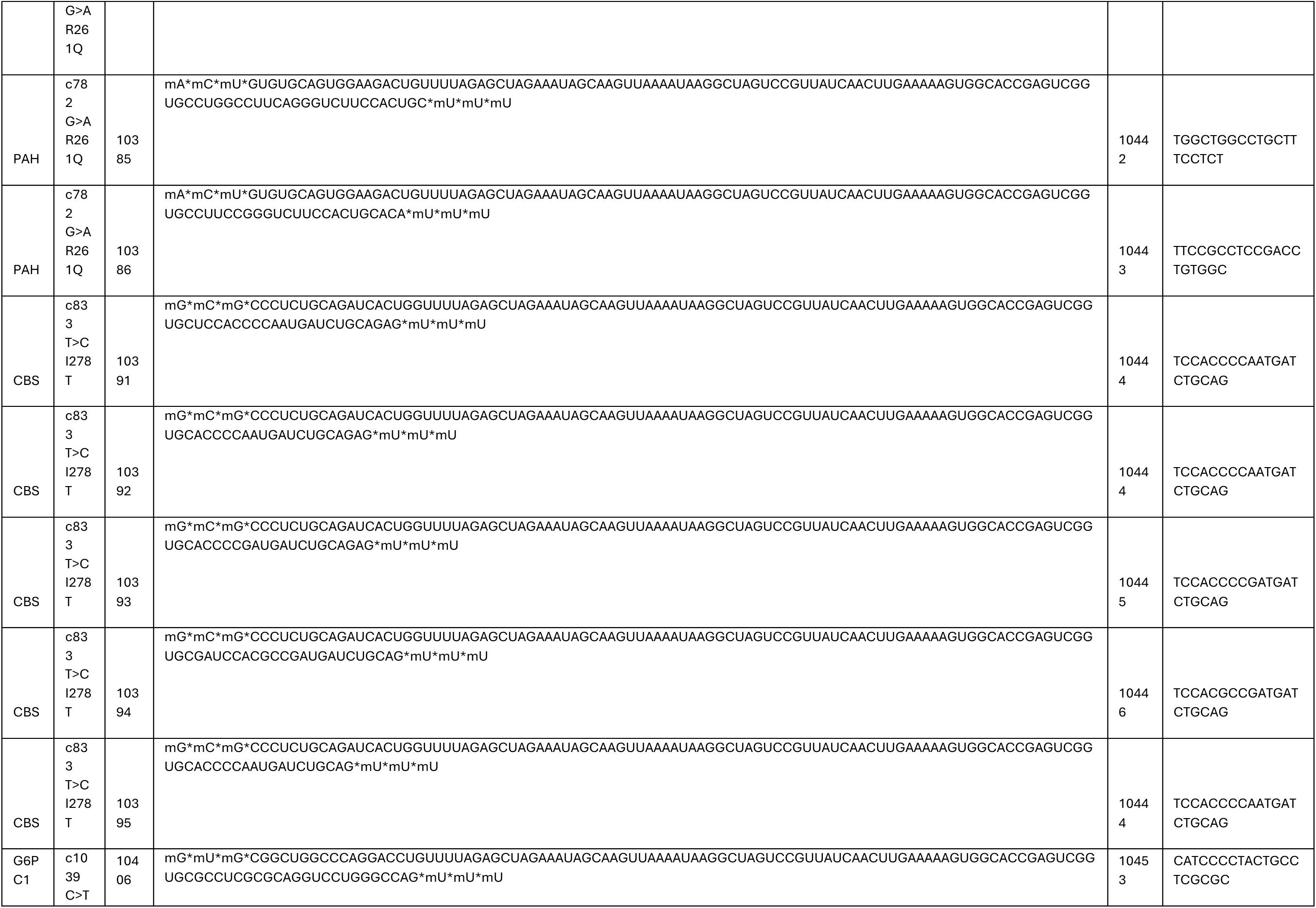
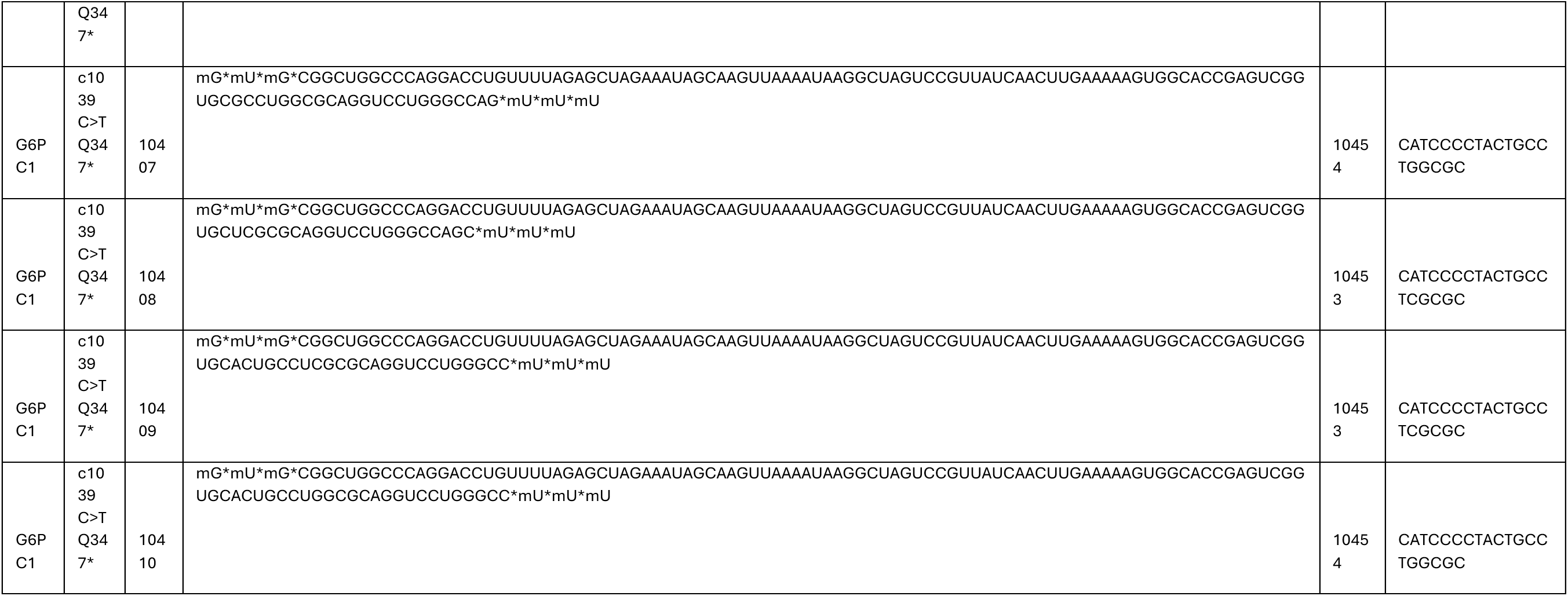
Sequences of sgRNAs and pegRNAs targeting mutations other than PAH p.R408W.

**Table IV.**
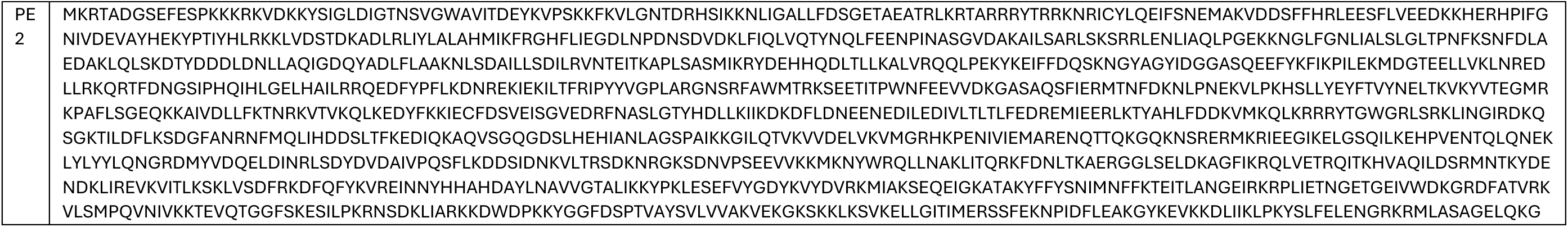
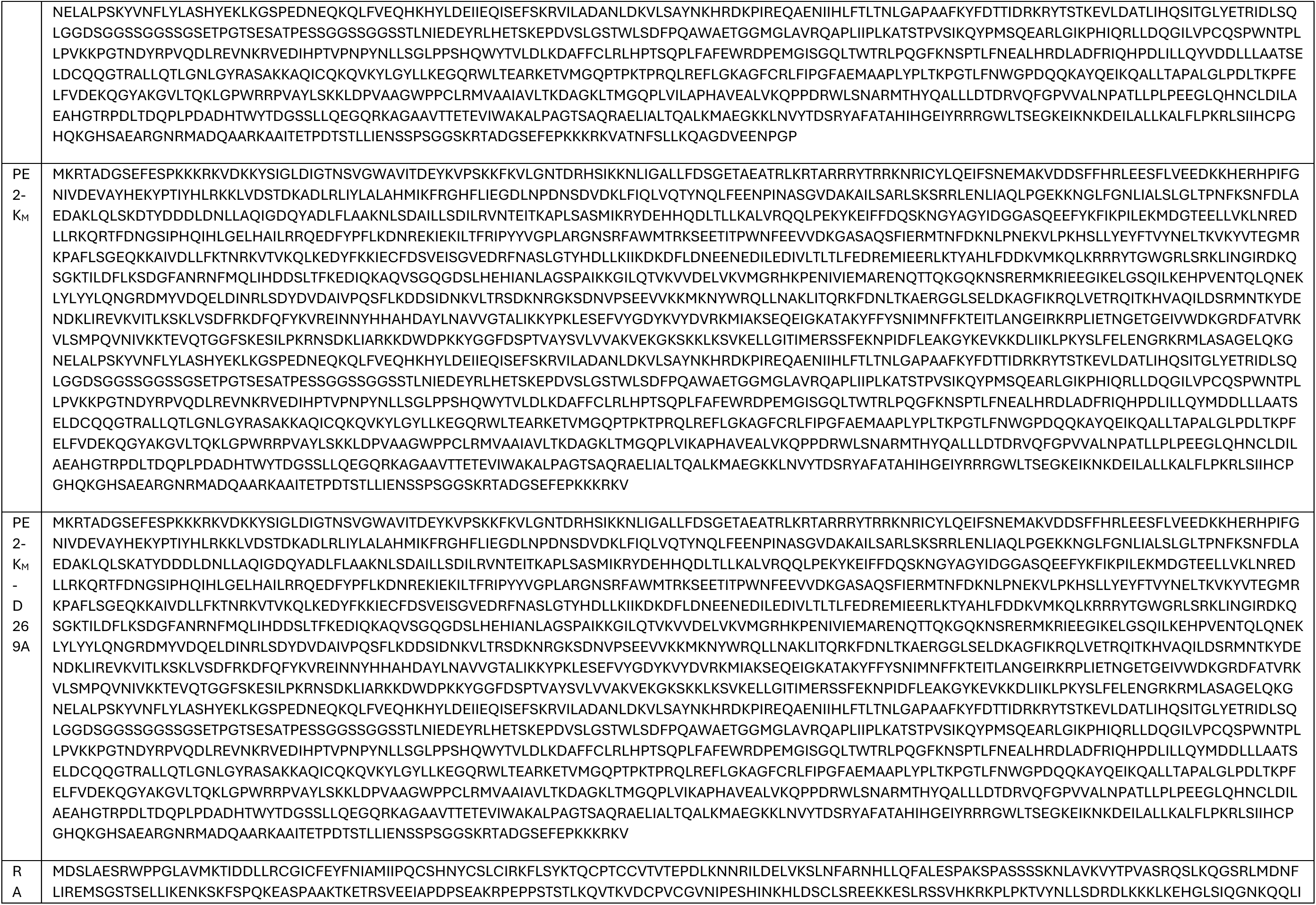
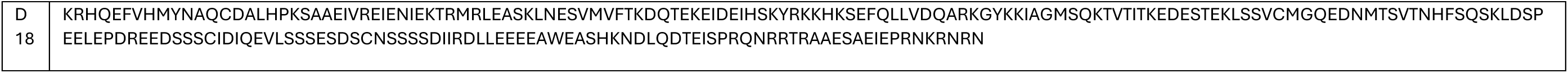
Sequences of gene editing proteins used in this study.

## Notes

### Competing Interest Statement

All Authors were/are paid employees of Moderna

